# Topological data analysis of pattern formation of human induced pluripotent stem cell colonies

**DOI:** 10.1101/2024.05.07.592985

**Authors:** Iryna Hartsock, Eunbi Park, Jack Toppen, Peter Bubenik, Elena S. Dimitrova, Melissa L. Kemp, Daniel A. Cruz

**Author notes:** these authors contributed equally to this work (Iryna Hartsock, Eunbi Park).

## Abstract

Understanding the multicellular organization of stem cells is vital for determining the mechanisms that coordinate cell fate decision-making during differentiation; these mechanisms range from neighbor-to-neighbor communication to tissue-level biochemical gradients. Current methods for quantifying multicellular patterning cannot capture the spatial properties of cell colonies across all scales and typically rely on human annotation or *a priori* selection of parameters. We present a computational pipeline that utilizes topological data analysis to generate quantitative, multiscale descriptors which capture the shape of data extracted from multichannel microscopy images. By applying our pipeline to certain stem cell colonies, we detected subtle differences in patterning that reflect distinct biological markers and progressive stages of differentiation. These results yield insight into directed cellular movement and morphogen-mediated, neighbor-to-neighbor signaling. Because of its broad applicability to immunofluorescence microscopy images, our pipeline is well-positioned to serve as a general-purpose tool for the quantitative study of multicellular pattern formation.

## Introduction

Fluorescence microscopy is one of the most important and widely used tools for studying cell physiology^1,2^, from intracellular interactions to multicellular organization and beyond. The processes affecting multicellular pattern formation are typically studied with multichannel microscopy, with the detection of spatial features and dynamics often relying upon the visual inspection of several images. Quantitative methods have been developed for recognizing phenotypes at designated spatial scales, and improvements in individual cell identification and segmentation within images of tissues have enabled the ability to characterize patterning as a collective property of multiple cells. For example, deep learning methods have been used for classification to compare and track the spatiotemporal characteristics of each cell^3^ for the definition of tissue features^4^. A different approach to quantifying multicellular organization uses temporal-spatial signal logic to comparatively score the complexity of an image to a target pattern^5^. While these data-driven tools are effective in computing the difference between patterns, their outputs (e.g. latent variables, similarity scores) do not necessarily provide biologically interpretable information on intercellular interactions.

Graph-based methods have also been applied to derive quantitative descriptors of a multicellular construct such as an aggregate or colony^6–9^. With this strategy, each pattern is considered as an ensemble of network connections between neighboring cells. Neighborhood network features (e.g. path lengths or a number of like-cell clusters) are extracted from segmented, digitized microscopy images for statistical comparison across conditions^9^ or are subsequently evaluated with dimension reduction techniques to classify or compare spatiotemporal patterns with other images^7,8^. However, these tools require some amount of human annotation or a selection of network metrics *a priori* to learn the data or define the reduced dimension metrics. Similar approaches which consider pairwise connections and distances between subcellular structures may require less supervision; these techniques use distance distributions to measure the correlation between each cell in a microscopy image^10–12^. Though they quantify some morphological details, these methods share the same limitation as several of the ones mentioned previously in that they can only capture structural features at a specific scale. By not considering how these features evolve across scales, such methods fail to incorporate the effects that different degrees of noise may have on a multicellular network^13^.

As an alternative to methods mentioned above, we propose to use topological data analysis (TDA) to examine pattern formation. TDA is a fast-growing field which provides methods for summarizing the shape of complex data^14^ and has found useful applications in several fields, including cosmology^15^, material science^16^, neuroscience^17^, anomaly detection^18^, and *C. elegans* behavior^19^. Persistence homology, the central tool of TDA, tracks the appearance and disappearance of structural features (e.g. bounded empty regions) of data across different scales^20^. A persistence diagram is the output of persistence homology, and it is stable under various data perturbations^21^. One can map persistence diagrams to persistence landscapes^22^ to bridge TDA with other data analysis techniques. Persistence landscapes are shape descriptors that live in a vector space and can be used as inputs to machine learning algorithms. Persistence landscapes have unique averages and satisfy the Strong Law of Large Numbers and the Central Limit Theorem, which makes them suitable for statistical inference.

Recent work established the feasibility of studying emergent spatial properties in developmental biology by combining TDA with a clustering method to study zebrafish patterns across agent-based model simulations^23^. Since then, approaches employing TDA have been applied to profile spatial configurations of epithelial cells^24^ and to classify differences in bone microstructure^25^. Landscapes derived from the multiparameter persistence homology have been utilized to identify spatial patterns of both imaged and simulated immune cells^26^, highlighting the translatable power of TDA-based approaches. The four instances above show how TDA can give insight into understanding multicellular organization; however, the tools and analyses are tailored to each case. Newly published work on efficient and automated multicellular patterns identification^27^ holds promise for broad applicability but currently relies on its application to simulated data. In^28^, the authors incorporated persistence landscapes into a microscopy image analysis pipeline (TDAExplore) for detecting changes in the architecture of actin cytoskeleton. While this tool is accessible and broadly applicable, it is limited to microscopy images with a single channel.

Here, we develop a computational pipeline for quantifying multicellular patterns observed in multichannel microscopy images using TDA. Three sequential modules form a complete pipeline which automates cell segmentation, cell type identification, and the generation of multiscale, topological descriptors (persistence landscapes) for a given microscopy image set. The TDA module outputs persistence landscapes of an image set which allows them to be combined with statistical and machine learning tools; the module also generates cycle representatives of detected structural features for each input image. The end result is a modular, general-purpose pipeline aimed at broadening the access to TDA in the context of microscopy images analysis.

We apply our pipeline to study stem cell colonies actively undergoing differentiation. In particular, we study differentiation in the context of human induced pluripotent stem cells (hiPSCs), which are reprogrammed from somatic cells and have the capacity to differentiate into all primary germ layers represented in embryos^29^. Because of their comparable characteristics to human embryonic stem cells^30^, hiPSC cultures have become powerful, patient-specific *in vitro* test beds for investigating the early stages of human embryonic development^31^. During differentiation, a population of hiPSCs undergoes multicellular self-organization which depends on intrinsic, autonomous properties of cells and intercellular communication involving molecules known as morphogens^32–35^.

To facilitate our study of hiPSC differentiation, we employ a cell line for which differentiation can be induced synthetically^36^. Prior imaging observations on this cell line imply that cell fate acquisition and localization are influenced by neighboring cells^36,37^. By applying our pipeline, we find that varying the amount of chemical induction reveals trends in pattern formation across stages of differentiation. We also show that studying spatial information enhances our ability to detect and examine these trends quantitatively. Finally, we analyze the patterning dissimilarities between all differentiated cells and those whose differentiation is synthetically induced. By doing so, we are able to draw a connection between local changes in the neighborhood composition of differentiated cells and system-level effects of synthetic induction. Thus, we provide evidence for how the interpretation of pattern formation can inform the discovery and study of the molecular mechanisms driving cell migration and cell fate decision-making.

## 1 Results

### Pipeline for Microscopy Image Analysis

Our main methodological contribution is a computational pipeline for microscopy image analysis that produces topological, multiscale descriptors of multicellular organization. Our goal in developing this computational framework is to open new avenues for the spatial cell analysis by making the tools from TDA accessible to a broader audience. To this end, we organized pipeline into three sequential modules: segmentation, cell type identification, and TDA. Each module is automated, can be applied to data from entire image sets, relies on a few user-set parameters if any, and requires little to no *a priori* knowledge to be used; see **Fig**. 1. First, the segmentation module obtains cell-specific locations and signal intensities of *n* biomarkers from each immunofluorescence microscopy image in the input set (**Fig**. 1**a**). Then, the cell type identification module categorizes each cell based on their signal intensities into one of 2^*n*^ cell types based on a user-selected percentile threshold (**Fig**. 1**b**). Finally, the TDA module derives topological descriptors of patterning for a given combination of the 2^*n*^ cell types (**Fig**. 1**c**-**h**).

**Figure 1.**
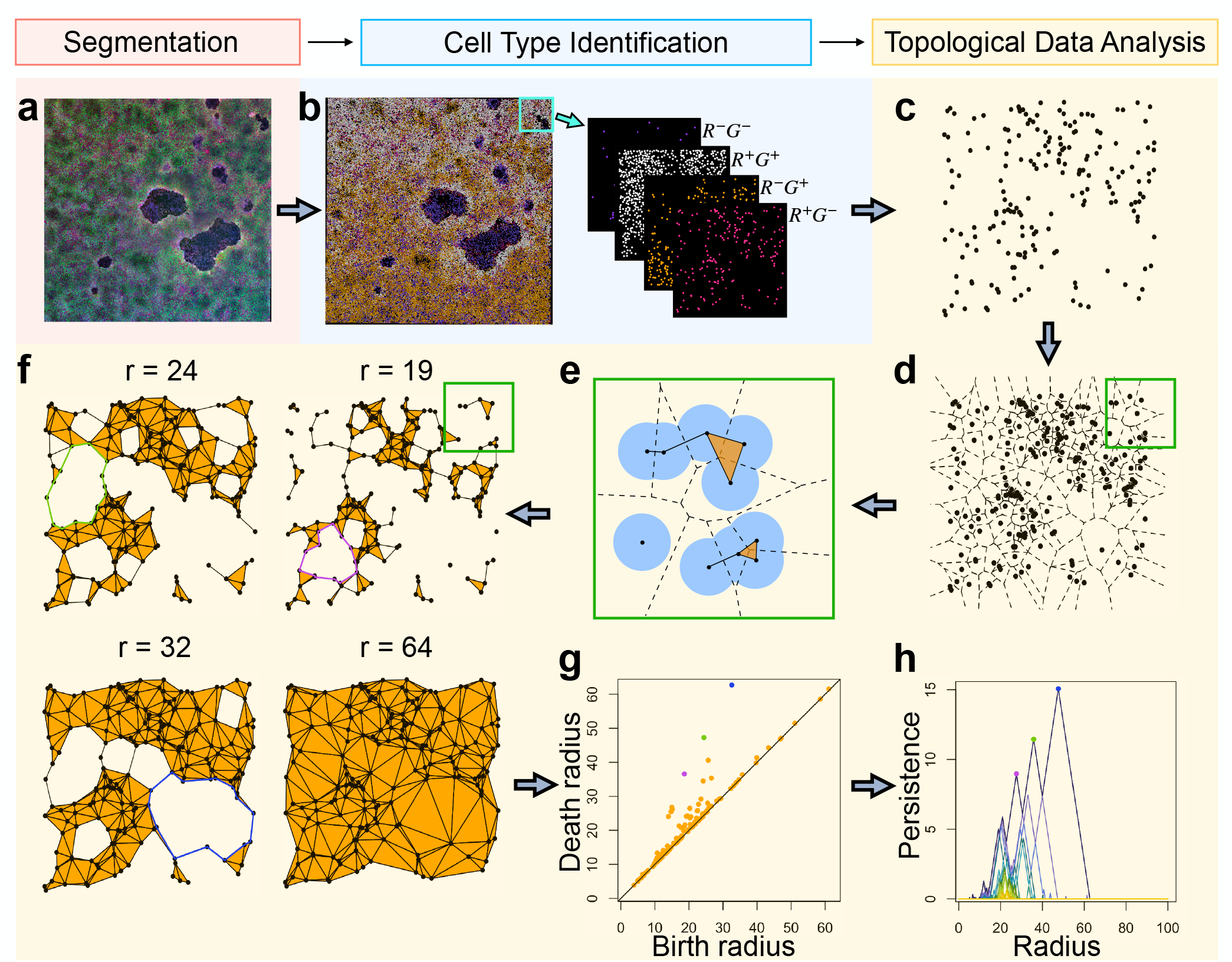
Our pipeline extracts topological descriptors from microscopy images of multicellular colonies. **a** An input microscopy image of cells with a nucleus signal in blue and two other signals: red (R) and green (G). This image is processed through the segmentation module of our pipeline to identify cell locations and associate a signal intensity to each cell. **b** A discretized version of the microscopy image in which cells are represented as points in the Euclidean plane and categorized into one of four cell types based on signal intensities by the cell type identification module. For the upper right patch, points in each cell type are shown. **c** The points of a single cell type (R^+^G^*−*^) in the patch. **d** The corresponding Voronoi diagram, a partition of the patch into regions enclosing a portion of the plane that is closest to each point. **e** A step in the Delaunay filtration for the top right part of the Voronoi diagram, built by connecting neighboring points with line segments and triangles using a proximity radius *r*. **f** Four stages in the Delaunay filtration of the patch, an increasing sequence of simplicial complexes, with the three most persistent enclosed empty regions shown at the radius *r* (in pixels) at which they appear. **g** The resulting persistence diagram, a topological summary that encodes the radii at which holes appear and disappear. **h** Its corresponding persistence landscape, a decreasing sequence of piecewise-linear functions, which can be combined with statistical and machine learning tools. This is the primary output of the TDA module in our pipeline.

The the three modules in our pipeline can be used in sequence to obtain topological descriptors from microscopy image sets without requiring users to perform additional computations like preprocessing their imaging data. Nonetheless, we have designed this computational framework to be modular so that users may swap one module with a similar tool of their choice as long as they respect the data formatting constraints of the next module in the sequence. The reason for this design is that users may prefer to employ tools which are more appropriate to their context than the general-purpose modules which we have written. In the case of the segmentation, we chose a histogram-thresholding based approach^38^ because it is one of the most straightforward and widely used methods for image segmentation^39^. There are several segmentation and image processing tools available^39–42^, including segmentation alternatives which extract information from microscopy images differently^43,44^. The approach we take to identify cell types is only based on signal intensities; see *Methods* for details. As such, it is broadly applicable because it does not rely on biological assumptions associated to a particular data set. Users may opt to identify cell types differently based on additional data or insights; there are even segmentation-free approaches to cell type identification^45^ that could be adapted to work with our pipeline. Lastly, there are several statistical approaches which have been traditionally applied to the context of analyzing spatial data^46,47^; these approaches may be used in addition to our TDA module for comparison and/or validation.

In the remainder of this section, we describe the basic principle of persistence homology^20^ which is employed in the last module of our pipeline and further detailed in the *Methods*. Let *P* be a collection of points on the plane (**Fig**. 1**c**). A *Voronoi cell* of a point *p* ∈ *P* is the region of the plane such that every point in that region is at least as close to *p* as to any other point in *P*.

The union of Voronoi cells covers the whole plane and is called a *Voronoi diagram* of *P* (**Fig**. 1**d**). Next, we gradually connect the points in *P* in the following way. First, we grow balls of radius *r* around every point in *P*. For every *p* ∈ *P*, we denote the intersection of its Voronoi cell and its *r*-ball by *V*_*r*_(*p*). For distinct *x, y* ∈ *P*, if *V*_*r*_(*x*) intersects *V*_*r*_(*y*), we connect *x* and *y* with a line segment. If for some *z* ∈ *P, V*_*r*_(*x*), *V*_*r*_(*y*), and *V*_*r*_(*z*) have a nonempty intersection, then we connect them in a triangle. This process is shown in **Fig**. 1**e**. If no four points in *P* lie on the same circle, then there will be no quadruple intersections. At a fixed *r*, we obtain an object consisting of vertices, edges, and triangles called a *simplicial complex*. By allowing *r* to grow, we get an increasing sequence of simplicial complexes called the *Delaunay filtration* (**Fig**. 1**f**)^48^. To every element of the filtration, we apply homology in degree 1 with binary coefficients, a method from algebraic topology that detects (1-dimensional) holes, namely, sequences of edges arranged in a cycle (see the colored cycles in **Fig**. 1**f**).

Persistence homology records the “births” and “deaths” of empty regions surrounded by cycles as the scale parameter *r* increases. This information is encoded as a collection of (birth, death)-pairs on the upper-left half-plane of **Fig**. 1**g**, and the resulting plot is called a *persistence diagram*. The distance from the points in the persistence diagrams to the diagonal is proportional to their *persistence*, which is the difference between their birth and death values. In order to apply statistics and machine learning to persistence diagrams, we convert them to persistence landscapes^22,49^. A *persistence landscape* (PL) is a decreasing sequence of piece-wise linear functions (*λ*_*k*_) with slopes 1, *−*1, or 0 plotted on the ((birth+death)*/*2, (death *−* birth)*/*2)-plane of **Fig**. 1**h**. The parameter *k* is referred to as the *depth* of the PL, with the first depth corresponding to the outermost function; see *Methods* for a precise definition of a PL. Unlike persistence diagrams, PLs have unique averages. We apply our pipeline to a small patch of hiPSC colonies in **Fig**. 1**c**-**h**. We have highlighted the three most persistence points in the persistence diagram (**Fig**. 1**g**), their corresponding representative cycles (**Fig**. 1**f**), and the corresponding peaks in the PL (**Fig**. 1**h**).

### Microscopy Imaging of Human Induced Pluripotent Stem Cells

We apply our pipeline to study differentiation and pattern formation in a genetically engineered hiPSC line capable of overexpressing transcription factor GATA6, a marker for differentiation potential^50^. We focus on the initial loss of pluripotency and thus consider protein markers GATA6 (differentiation) and NANOG (pluripotency). This cell line has been transduced with a gene construct (**Fig**. 2**a**) to express HA epitope-tagged GATA6 (GATA6-HA) transgenes by Doxycycline (Dox) induction, which results in rapid colony patterning between pluripotent GATA6cells and GATA6+ cells^36^. The extent of the GATA6-HA activation can be regulated by the amount and duration of the Dox treatment. We tested and imaged different Dox concentrations of 0, 5, 15, and 25 ng/ml for three days (**Fig**. 2**b** and **Supplementary Fig**. S1). All microscopy images were generated by 6 *×* 6 multiple fields combined to yield spatial patterns in a large scope of view so that the average image size is 2, 850 *×* 2, 850 pixel (4, 132.5 *×* 4, 132.5*µ*m); see *Methods* for the pixel conversion. We observed that higher Dox concentration resulted in increased pan-GATA6 and HA expression levels, and that pan-GATA6 expression was likely to be higher than that of HA (**Fig**. 2**b**-**c**).

**Figure 2.**
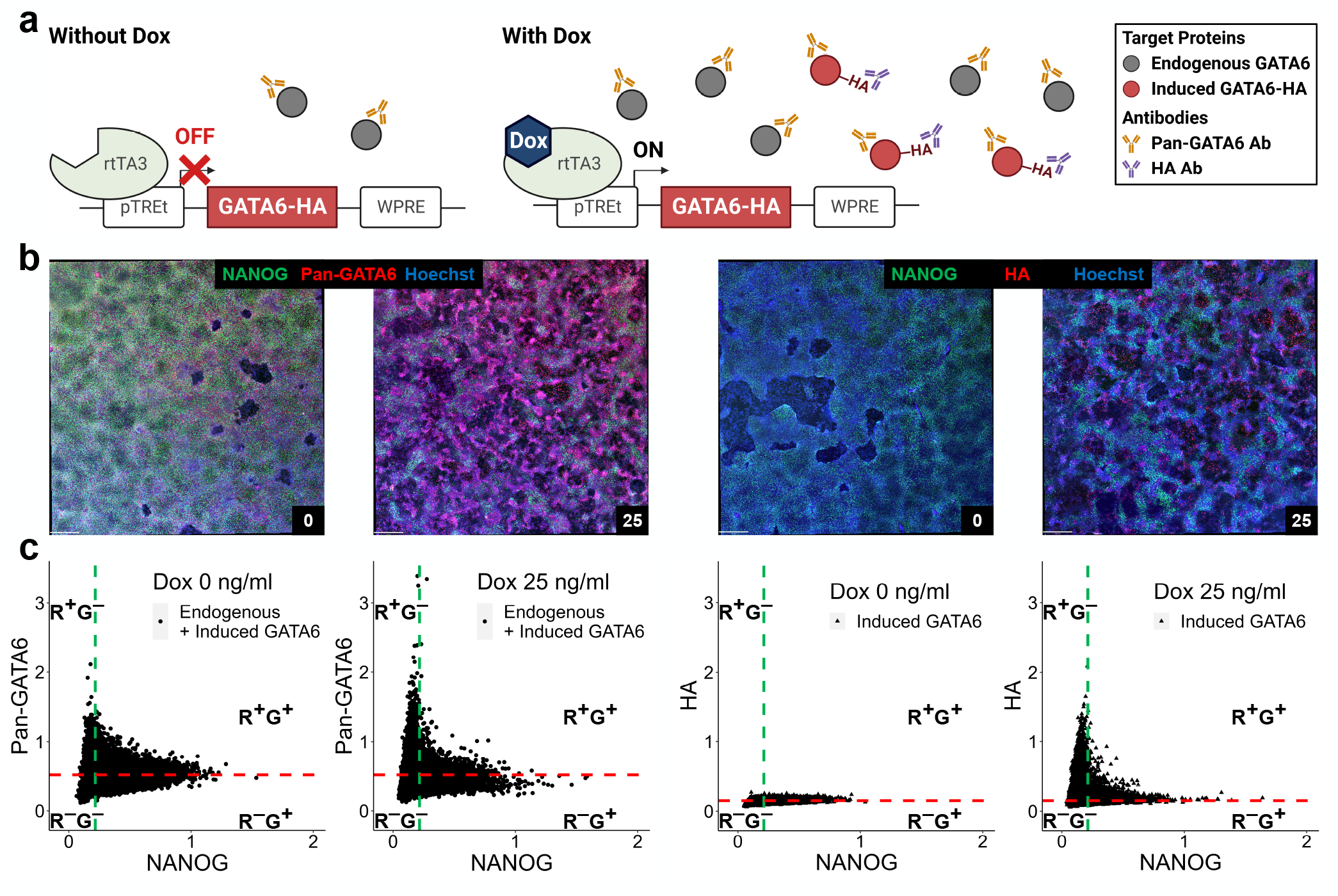
Artificial induction of exogenous GATA6-HA occurs within the context of endogenous GATA6 expression. **a** Gene circuit for chemical induction of GATA6-HA expression. The pan-GATA6 antibody can detect both endogenous GATA6 and induced GATA6-HA, whereas the HA antibody can only detect induced GATA6-HA. **b** Representative immunofluorescence images of NANOG and pan-GATA6 or HA at 0 and 25 ng/ml Dox concentrations (scale bar, 440*µm*). Using Volocity, we applied the gamma changes (gamma of 1.5) after brightness enhancement on all stitched large images used in the figure for better contrast for representation (see *Methods*). **c** Quantification of segmented images by each channel from **b**. Green (NANOG) and red (pan-GATA6 or HA) fluorescent intensities are normalized to the corresponding nuclear Hoechst value. The threshold for the green channel is given by the green dotted line, and the red dotted line indicates the threshold for the red channel. The four cell types based on these thresholds are given by the labels R^+^G^+^, R^+^G^*−*^, R^*−*^G^*−*^, and R^*−*^G^+^ of the corresponding quadrant.

Pan-GATA6 quantification represents summative GATA6 expression levels in each cell of endogenous GATA6 and induced GATA6, whereas the HA levels only represent the GATA6-HA expression induced by the gene circuit.

Using our pipeline, we segmented the images based on nuclear morphologies and acquired cellular positional information and intensity profiles of the immunofluorescent markers. The green fluorescent signal serves as a marker of NANOG expression, and the red fluorescent signal corresponds to either immunolabeling of pan-GATA6 or HA in the corresponding cell cultures. Every population of cells was discretized into binary groups using threshold values for each channel intensity. Due to computational limitations, we split each discretized image into 16 even, non-overlapping patches. As the result, in each Dox concentration group we have 240 patches (from 15 images) separately in pan-GATA6 and HA populations. We provide full details on our thresholding in the *Methods* and give an idea of our approach below for a given population of cells. The value corresponding to the 75^th^ percentile of the green intensity distribution for the *highest* Dox treatment group (25 ng/ml) was assigned as the green threshold for all Dox treatment groups. All cells with green signals above this threshold are categorized as green positive, G^+^ (**Fig**. 2**c**). On the other hand, the values corresponding to the 75^th^ percentile for the *lowest* Dox treatment groups (0 ng/ml) were assigned as the red (pan-GATA6 or HA) thresholds for all Dox treatment groups. All cells in the pan-GATA6 or HA populations with red signals above their corresponding red threshold are categorized as red positive, R^+^ (**Fig**. 2**c**). We selected the 75^th^ percentile after assessing how the populations were partitioned into each cell type for different percentile choices (**Supplementary Table** S1); see *Methods* for details.

To verify that the pattern formation observed in the Dox treatment groups (**Fig**. 2 and **Supplementary Fig**. S1) does not arise prior to the treatment protocol, we also obtained microscopy images at the start of the protocol (**Supplementary Fig**. S2) and processed them with our pipeline as well. We make the simplifying assumption that pluripotent cells are R^*−*^G^+^ cells in the treatment groups because GATA6 is a marker for differentiation potential^50^ and because NANOG is a marker for pluripotency^51–54^. It follows that R^*−*^G^+^ cells are likely the most biologically similar to the (pluripotent) cells in the initial condition group (i.e. those present at the start of the protocol).

### Quantifying Differences in Pattern Formation Induced by Varying Doxycycline Concentration

In **Fig**. 2 and **Supplementary Fig**. S1, we notice clear differences in the spatial organization of cells across the Dox treatment groups for both the pan-GATA6 and HA populations; we also notice differences between these groups and the initial condition group (**Supplementary Fig**. S2). We conducted two-sample permutation tests on the PL vectors from the initial condition group and the PL vectors of R^*−*^G^+^ cells from each Dox treatment group in pan-GATA6 and HA populations; see *Methods* for more details. For each Dox treatment group in the pan-GATA6 and HA populations, the p-value of each permutation test was below 5E*−*2, a statistical significance threshold used in this work (**Supplementary Table** S2). This indicates that the cell patterning structure in all Dox treatment groups differs from the one in the initial condition group. The histograms of the R^*−*^G^+^ cell counts in the Dox treatment groups and total cell counts in the initial condition group are shown in **Supplementary Fig**. S9 and **Supplementary Fig**. S10**a**, respectively.

To study the Dox treatment groups further, we generated the average PL of each cell type (R^+^G^+^, R^+^G^*−*^, R^*−*^G^*−*^, and R^*−*^G^+^) in the pan-GATA6 (**Supplementary Fig**. S3) and HA (**Fig**. 3 and **Supplementary Fig**. S4) populations per treatment group. For reference, the average PL of the initial condition group is shown in **Supplementary Fig**. S10**b**. We also ran permutation tests on PL vectors of each pair of Dox concentrations (per cell type). The p-values for almost all pairwise permutation tests are less than 5E*−*2 (**Supplementary Table** S3), substantiating the idea that topological features differ from one Dox concentration to another for both pan-GATA6 and HA groups. The only p-values greater than 5E*−*2 are associated to the R^*−*^G^*−*^ cell type in the pan-GATA6 group, which implies that pattern structure within this cell type does not change much across Dox concentrations. This observation is also supported by the visual similarity of their average PLs (**Supplementary Fig**. S3). Using our 75^th^ percentile thresholds (**Fig**. 2**c**), we organize our analysis of the average PLs in **Supplementary Fig**. S3, **Fig**. 3, and **Supplementary Fig**. S4 based on R^+^ and R^*−*^ cell populations.

**Figure 3.**
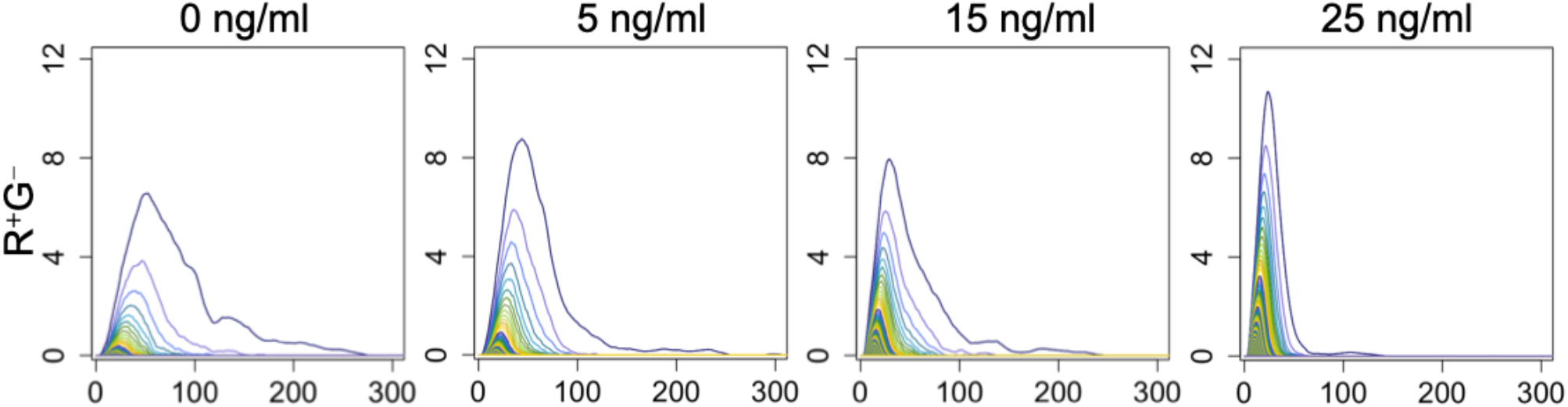
Comparison of average persistence landscapes across various Dox concentrations for R^+^G^*−*^ cell type in the HA group. As Dox concentration increases, the average persistence landscape gets taller and narrower.

We observe trends that correspond to an increase in differentiation from the population distributions (**Supplementary Table** S1) and the average PLs of R^+^ cells in the pan-GATA6 (**Supplementary Fig**. S3) and HA (**Fig**. 3 and **Supplementary Fig**. S4) groups. As expected, the number of R^+^ cells increases and the number of R^*−*^ decreases as the Dox concentration increases. The average PLs of R^+^G^*−*^ cells are compressed left, indicating that most cycles in higher Dox treatment groups are born earlier (**Supplementary Fig**. S3 and **Fig**. 3). This trend suggests that differentiated cells stay in close proximity to one another even as their number increases. The increasing height of average PLs (across all depths) also demonstrates that the cycles in higher Dox treatment groups are also more persistent, with labyrinthine patterns emerging in the 25 ng/ml group. At the first depth (the outermost function), the height of the average PLs of R^+^G^+^ cells also increases with Dox concentration in the pan-GATA6 and HA groups; however, after the first depth, the nested heights of the averages PLs do not always follow this trend.

In general, the trends of R^*−*^G^+^ cells in the pan-GATA6 (**Supplementary Fig**. S3) and HA (**Fig**. 3) groups are reversed from those of R^+^G^*−*^ cells. Average PL heights of R^*−*^G^+^ cells decrease for all depths in the HA population but only after the first few depths in the pan-GATA6 population. This behavior corresponds to the fact that most of the cycles in R^*−*^G^+^ populations are less persistent in higher Dox treatment groups. The average PLs also widen as the Dox concentration increases, indicating that cycles in R^*−*^G^+^ populations have a wide range of births in higher Dox treatment groups. Together, these observations suggest that most pluripotent cells spread out from each other as their number decreases due to differentiation. The average PLs of R^*−*^G^*−*^ cells have the same behavior as the average PLs of R^*−*^G^+^ cells in the HA group at most depths (**Supplementary Fig**. S4); however, the average PLs in the pan-GATA6 population look similar across most Dox treatment groups (**Supplementary Fig**. S3).

### Spatial Information Improves Classification and Prediction of Doxycycline Treatment Groups

To determine if the variations of the topological signatures produced by our pipeline are sufficient to classify image patches according to their Dox concentration, we applied multiclass support vector machines (SVM) to concatenations of vectors generated from the PLs of all cell types (per patch). For comparison, we also performed the same analyses on cell count data by constructing a four-dimensional vector for each patch which holds the count for each cell type. Note that we normalize both PL and cell count vectors; see *Methods* for more details about SVM. We provide the confusion matrices of one instance of multiclass SVM in **Supplementary Table** S4 for the PL vectors and cell count vectors. Averaging across 20 instances of multiclass SVM, we accurately classified 71.8% patches in the pan-GATA6 group and 73.5% in the HA group with their corresponding Dox concentrations using PLs. The average accuracy dropped to 67% for the pan-GATA6 group and 68.3% for the HA group when we used the cell count vectors instead, indicating that including spatial information improves our ability to distinguish patches between various Dox concentrations.

Since the Dox concentration is a numerical variable, we also performed support vector regression (SVR) to predict the Dox concentration of each image. We conducted SVR separately on the normalized PL vectors and normalized cell count vectors of each image’s constituent patches; see *Methods* for details. We then averaged the Dox concentration predictions of an image’s patches. The resulting Dox concentration predictions are shown in **Fig**. 4 using PL vectors and in **Supplementary Fig**. S5 using cell count vectors. **Supplementary Table** S5 has additional information on every box plot (**Fig**. 4 and **Supplementary Fig**. S5), including the deviation of the median from the actual Dox concentration (called “error”). By comparing the two approaches, we can observe that the errors are higher overall for the cell count vectors than for the PL vectors. The Dox concentration predictions on PL vectors for 5 ng/ml, 15 ng/ml, and 25 ng/ml are well separated from one another while predictions on cell count vectors noticeably overlap (**Fig**. 4 and **Supplementary Fig**. S5). In most Dox treatment groups, we also observe that the interquartile ranges of predictions are narrower and the whiskers are shorter using PL vectors instead of cell count vectors.

**Figure 4.**
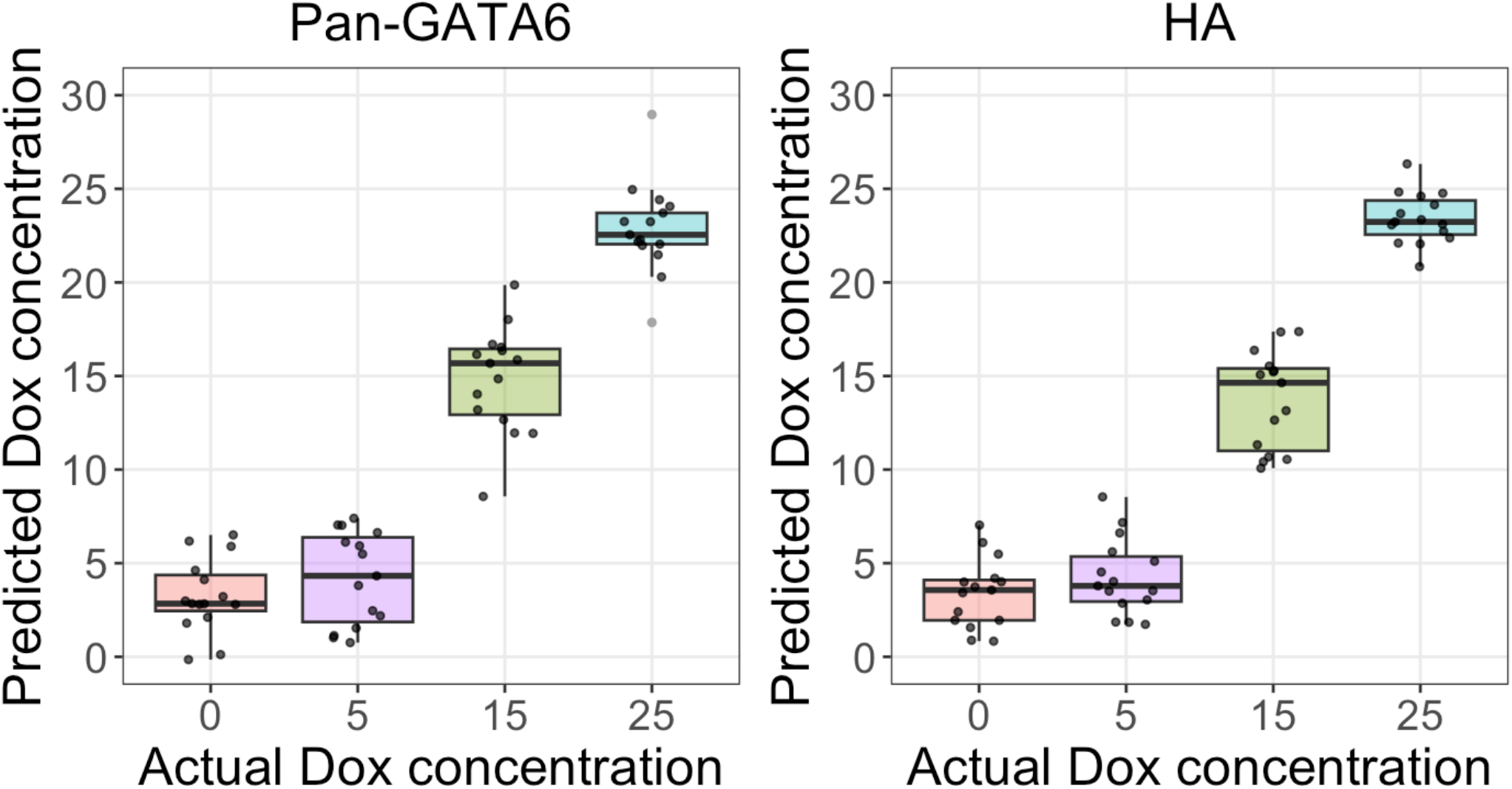
We performed support vector regression on the persistence landscape vectors extracted from patches; we then averaged the Dox concentration predictions of each image’s patches. The image predictions of each Dox treatment group are spread horizontally for better visualization, and the outliers are marked with gray points. The Dox concentration predictions of 5 ng/ml, 15 ng/ml, and 25 ng/ml images are close to their actual values and well separated from one another, but the Dox concentration predictions of 0 ng/ml images are not well distinguished from 5 ng/ml images.

By considering **Supplementary Table** S4 and **Fig**. 4, we notice that it is difficult to distinguish between 0 ng/ml and 5 ng/ml Dox concentrations with SVM and SVR in both pan-GATA6 and HA populations. This suggests that the 5 ng/ml increase in Dox concentration does not accelerate the differentiation of cells enough to change the pattern formation so that the SVM/SVR models can properly learn the difference between these Dox treatment groups. Although the permutation test on 0 ng/ml and 5 ng/ml Dox treatment groups shows that their topological structures are distinct (**Supplementary Table** S3), the ranges of their Dox concentration predictions greatly overlap (**Fig**. 4). Multiclass SVM and SVR on cell count vectors also do not separate well between 0 ng/ml and 5 ng/ml Dox treatment groups (**Supplementary Table** S4 and **Supplementary Fig**. S5). Because most incorrect classifications in the multiclass SVM confusion matrices involve 0 ng/ml and 5 ng/ml (**Supplementary Table** S4), we performed pairwise SVM on the PL and cell count vectors of two distinct Dox treatment groups; see *Methods* for more details. Pairwise SVM accuracy is high (i.e. greater than *∼* 90%) for almost every Dox treatment group pair using PL vectors (**Supplementary Table** S6); only the 0 ng/ml vs 5 ng/ml classification had a low accuracy (61.81% for the pan-GATA6 group and 62.69% for the HA group). Pairwise SVM accuracies on the cell count vectors are lower than those on the PL vectors, except for the 0 ng/ml vs 5 ng/ml classification which is nevertheless fairly low.

### Distinguishing Pattern Formation of Total GATA6 vs Synthetically Induced GATA6

We consider the following question: “Does the choice of a biological marker, pan-GATA6 or HA, affect how we perceive the structure of differentiated cell colonies?” Recall that the HA-specific antibody only targets synthetically induced GATA6 expression whereas the pan-GATA6 antibody stains for total GATA6 expression (detecting both endogenous GATA6 plus synthetically-induced GATA6). Because prior work suggests the local expression of GATA6 in neighboring cells may result migration and cell fate specification^36,37^, addressing this question could imply the presence of intercellular communication among neighborhoods of cells expressing (endogenous and/or synthetic) GATA6. To address the question above, we examine cells with high red (pan-GATA6 or HA) and low green (NANOG) intensity in the highest (25 ng/ml) Dox treatment group. We observed subtle, visual differences (e.g. number and size of holes) between microscopy images from both pan-GATA6 and HA populations (**Fig**. 2**b** and **Supplementary Fig**. S6). However, our goal is to quantitatively determine if there is a difference in the pattern formation of these two populations.

We performed a permutation test on PL vectors from the pan-GATA6 and HA populations to determine if the cell differentiation patterns are statistically distinct; see *Methods* for details. We obtained a p-value of 1E*−*4 which strongly supports the claim that the topological structure of these patterns are different. For comparison, we computed the local density of R^+^G^*−*^ cells in the pan-GATA6 and HA groups. This difference was statistically significant (p-value less than 2.2E*−*16) when using all density values due to the large sample sizes of each group; however, when we used the mean density per image, the p-value rose to 5.605E*−*2. We also performed k-means clustering with the mean local density and intensity profiles, which did not yield a clear partition between the pan-GATA6 and HA groups (**Supplementary Fig**. S7). Thus, we find that these traditional approaches to spatial statistics are not sensitive enough to detect the subtle differences between the two populations. In particular, we observe an advantage to having a multiscale, topological representation (PL) of the patterns in these population over one measurement (local density) at a fixed scale.

Our next goal is to understand the dissimilarities between the cell differentiation patterns of pan-GATA6 and HA groups; see **Supplementary Fig**. S6 for example. We can visualize these dissimilarities by subtracting the HA average PL (**Fig**. 5**a** middle) from the pan-GATA6 average PL (**Fig**. 5**a** left) and plotting the difference (**Fig**. 5**a** right). We observe two distinct parts in the difference plot separated by the vertical (gray) line defined by the equation (birth+death)*/*2 = 35. We begin by observing that the side left of the vertical line is negative (**Fig**. 5**a**), indicating that the cycles that appear early in the HA group are more persistent than those in the pan-GATA6 group. Cycles that are more persistent surround larger empty regions, so HA populations have more of these regions than pan-GATA6 populations. This difference can be explained by the fact that HA antibody only detects induced GATA6 expression; thus, neighboring R^+^G^*−*^ cells in the HA populations may be kept from entering empty regions by other cells with high levels of total GATA6 expression.

**Figure 5.**
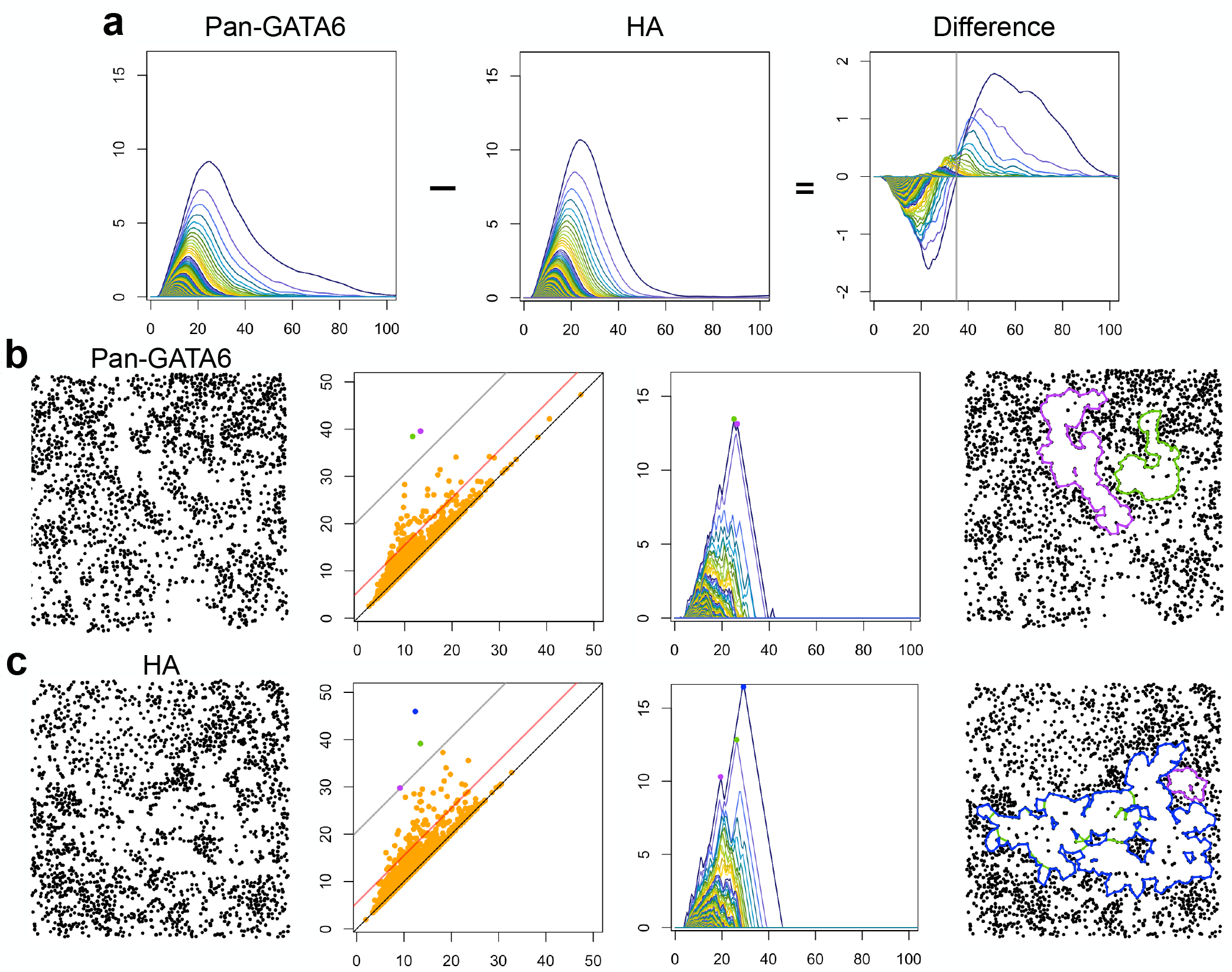
Comparison between pattern formations in the pan-GATA6 and HA groups for R^+^G^*−*^ cell type with 25 ng/ml Dox concentration. **a** Average persistence landscapes of the pan-GATA6 and HA groups and their difference. The gray vertical line splits the difference plot into two parts showing distinct types of dissimilarities between the cell differentiation patterns. **b** The TDA pipeline applied to a representative patch of the pan-GATA6 group, and **c** a representative patch of the HA group. For each patch, its persistence diagram, persistence landscape, and most persistent cycles are shown. In the persistence diagrams, the gray line indicates the persistence threshold of 20 and the red line marks the threshold of 5. There are 2 points above the gray line in the pan-GATA6 patch and 3 points in the HA patch; the corresponding representative cycles of these points are shown in the rightmost column. Note that the green cycle inside the blue one in the HA patch.

To better grasp the difference in number of large empty regions in the pan-GATA6 and HA groups, we analyze the points in their persistence diagrams. In particular, we count the points are above certain persistence thresholds and satisfy the inequality (birth+death)*/*2 *<* 35, which corresponds to the left side of the difference plot (**Fig**. 5**a**). We choose low (5) and high (20) thresholds to identify points corresponding to medium and large holes without considering the points that capture biological noise. Fixing the persistence threshold at 20, there are 182 more of these points in the HA group than in the pan-GATA6 group; the average number of such points is 3 in HA and 2 in pan-GATA6. If we lower the persistence threshold to 5, then the difference in number increases to 2368, and the averages become 84 for HA and 74 for pan-GATA6. We choose two representative patches from each group with about 1, 900 cells each to visualize these dissimilarities (**Fig**. 5**b**-**c**).

Now, we focus on the right side of the difference plot (**Fig**. 5**a**), which is positive because the average PL of the pan-GATA6 group is wider than the average PL of the HA group. This implies there are more cycles that are born later in the pan-GATA6 populations than in the HA populations; most appear in patches from the pan-GATA6 group where cells enclosing large empty regions are relatively far apart. By analyzing the histograms of cell counts per patch of both groups (**Supplementary Fig**. S8), we observe that the count distribution of the pan-GATA6 group is also wider than that of the HA group. This observation provides further evidence for our assertion about the heterogeneous spread of R^+^G^*−*^ cells in the pan-GATA6 group. There are several patches in the pan-GATA6 group whose cell counts are below the count distribution of the HA group; most contain large empty regions which are enclosed by few cells. The existence of these patches results in the elevated region on the right side of the difference plot (**Fig**. 5**a**). If we omit patches with less than 1, 000 cells and re-compute the two average PLs for the remaining patches, then the elevated region mostly diminishes in the difference plot (**Supplementary Fig**. S11). Thus, the presence of large holes which appear later is not representative of the pan-GATA6 pattern formation.

In order to validate our findings, we conducted additional analysis using stitched image files without brightness and shading corrections (see *Methods*). For 25 ng/ml Dox treatment group, we computed the PLs for R^+^G^*−*^ cells from this approach in all patches belonging to both pan-GATA6 and HA groups. We performed a permutation test on the resulting PL vectors, yielding a p-value of 1E*−*4. This outcome reaffirms our earlier observation, emphasizing the inherent divergence in the topological structure of differentiation patterns between the pan-GATA6 and HA groups. Moreover, in the difference plot of the average PLs of the two populations (**Supplementary Fig**. S12), we observed a single large negative region similar to the one present in (**Fig**. 5**a**) and (**Supplementary Fig**. S11). We also quantified the average number of points in the persistence diagrams satisfying the condition (birth+death)*/*2 *<* 35 across all patches, with a persistence threshold of at least 20. This count remained consistent, with 2 points identified in the pan-GATA6 group and 3 in the HA group, mirroring our earlier observations.

This additional analysis strongly supports our earlier discovery about the differences between the HA and pan-GATA6 groups.

## Discussion

We introduced a general-purpose pipeline to quantify cell pattern formation from microscopy images and applied it to images of human induced pluripotent stem cell (hiPSC) colonies. Persistence homology, the main tool of topological data analysis (TDA), allowed us to capture holes in the spatial organization of cells at multiple scales. We used functional representations of persistence homology called persistence landscapes (PLs) to summarize and compare the multicellular patterns from various experimental conditions. By using our pipeline, we successfully distinguished features that reflect the degree of differentiation within an imaged colony as well as the subtle differences between immunofluorescence patterning generated by antibodies targeting a subset (HA) versus the total level (pan-GATA6) of a protein (GATA6). Our topological summaries revealed trends of how the spatial organization of each cell type changes as Dox concentration increases. In particular, our results suggest that differentiated cells remain near each another even as their number grows with increasing synthetic induction. On the other hand, most pluripotent cells seem to disperse from each other as their numbers shrink due to differentiation. SVM classification and SVR prediction on PL vectors of different Dox treatment groups yielded better accuracy than on simpler cell count vectors, indicating that multicellular spatial information is richer and more valuable for distinguishing between various Dox concentrations.

Our work also demonstrated that dissimilarities in the spatial patterning of the pan-GATA6 and HA-tagged populations are statistically significant. We were able to detect that the HA group, which reflects synthetic induction of GATA6, has more persistent cycles than the pan-GATA6 group using PLs; these cycles correspond to ample (relative to the distance between neighboring cells) empty regions within the microscopy images. Further investigation showed that the HA populations had more large empty regions than the pan-GATA6 populations. Because pan-GATA6 detects induced and endogenous GATA6 levels, the difference in patterning between these groups may be caused by heterogeneous expression of endogenous GATA6 in the pan-GATA6^high^NANOG^low^ populations and the migration of HA^high^NANOG^low^ cells towards each other because of neighbor-to-neighbor signaling in the context of morphogen diffusion. To investigate this matter further, we imaged cell populations co-stained with both pan-GATA6 and HA antibodies; see **Fig**. 6a. Combined with our quantitative analysis, these images suggest that holes in HA^high^NANOG^low^ cell populations may be partially occupied by other cells expressing (endogenous) GATA6, which would be detected with the pan-GATA6 marker.

**Figure 6.**
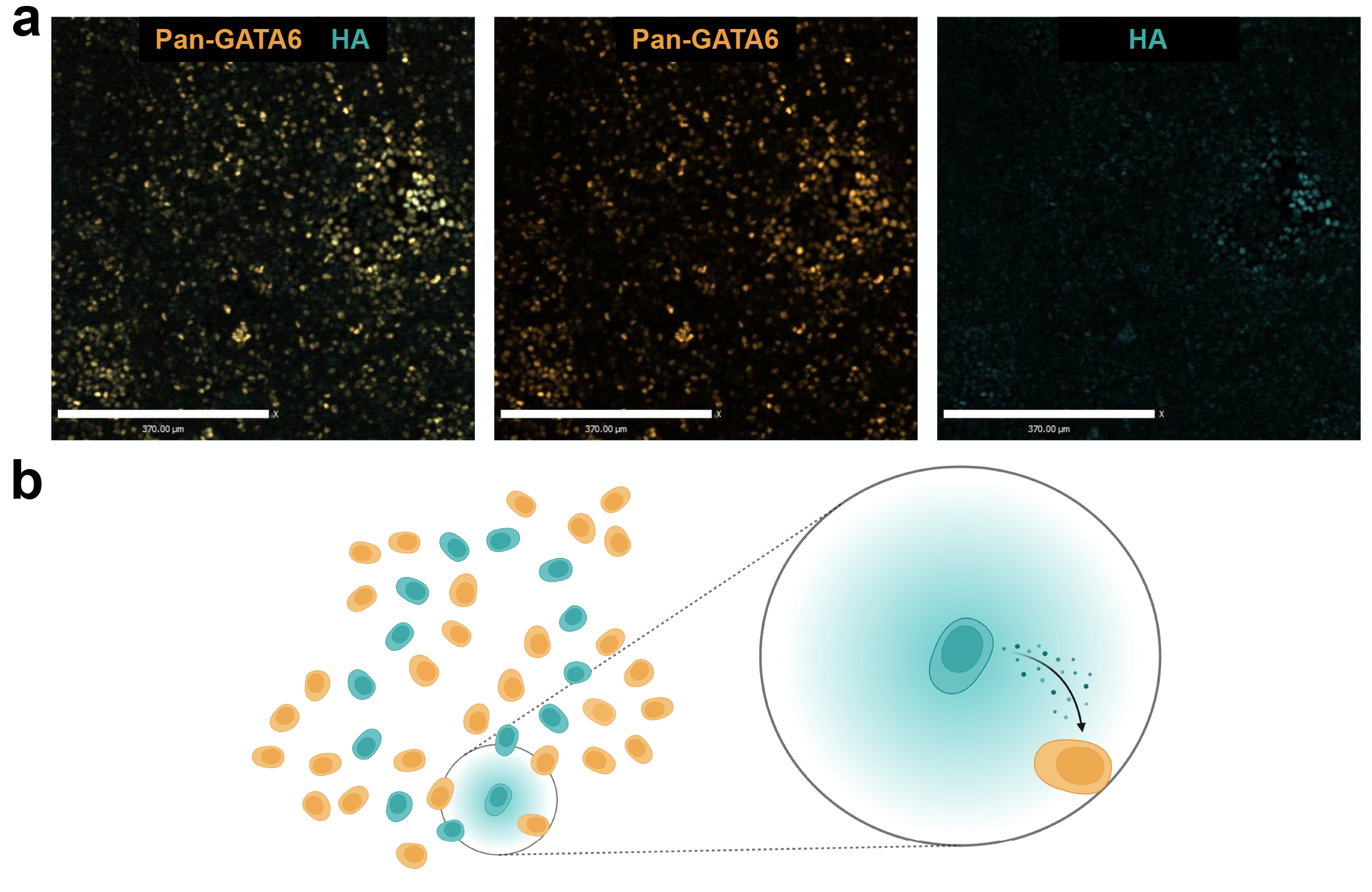
The topological structures of the pan-GATA6 and HA populations are statistically different according to our results. This difference may be due to cell migration and intracellular and/or intercellular communication in response to morphogen gradients. **a** Immunofluorescence images of co-stained pan-GATA6 and HA antibodies at 25 ng/ml Dox concentration (scale bar, 370*µ*m). The greater presence of cycles in the HA group which are more persistent suggests that cells whose differentiation is synthetically induced may be in close proximity due to chemotaxis. **b** A schematic representation of our proposed morphogen-mediated signaling, where heterogeneity caused by endogenous GATA6 levels may give rise to less persistent holes in the pan-GATA6 group.

Observations from^36,37^ about the hiPSC line used in this work indicate that the local expression of GATA6 in neighboring cells and/or the presence of neighboring differentiated cells influence cell fate acquisition and localization. It is possible that HA^high^NANOG^low^ cells remain relatively close to one another over the course of differentiation because of mitosis; however, it is not known whether this cell line undergoes symmetric or asymmetric cell division^55–58^. Alternatively, we propose that the difference in the number and persistence of cycles in the pan-GAT6 and HA populations implies the existence of cell migration and intracellular and/or intercellular communication in response to morphogen gradients (**Fig**. 6b). Further experimentation using time-course data would be required to determine if such preferential cell migration occurs and/or if morphogen gradients drive pattern formation. As a whole, our multi-scale analysis serves as an example of how measuring and interpreting topological features in multicellular pattern formation can inform the discovery and study of the molecular mechanisms driving cell development and behavior.

Our pipeline has several advantages, including its ability to capture the structure of cell patterns across all scales, but it has some limitations as well. The discretization process in the cell type identification module relies on the selection of a threshold; minor alterations of this parameter produce small changes in the populations of each cell type and thus may affect downstream analysis. We chose this threshold carefully by examining the population distributions of each cell type; see *Methods* for details. A benefit of TDA is that the persistence diagram and PL are insensitive to various imperfections in the microscopy images and their discretizations, including small changes in the positions of the cells and the misidentification of a pair of neighboring cells as a single cell, or vice versa. However, the persistence diagram and PL are sensitive to other changes, such as a cell with no nearby cells being omitted, being erroneously added, or being mislabeled due to a small change in its color intensity. Such color intensity changes may arise during image correction and/or be caused by stitching microscopy image tiles to produce large images (as done in our data set). There are emerging technologies which allow for microscopy-based imaging of a large field of view without the use of stitching^59,60^; our pipeline could be readily applied to such image outputs because of its general-purpose design. The sensitivity to changes in channel intensity near the channel threshold may be overcome with more computationally intensive TDA methods such as perturbation and averaging^61^ or multiparameter persistent homology^26^. The sensitivity to outliers may be overcome by subsampling^62^ or denoising^63^. In our approach, these shortcomings are alleviated by averaging over a large number of patches for each image.

The differentiation of stem cells is one of several contexts in which patterns arise based on interactions within a microenvironment. Other contexts include tumor architecture, bacterial biofilm formation, and particle organization on surface materials^64–67^. Research on spatial dynamics has been drastically growing within the past 5 years, primarily led by spatial transcriptomics^68^. On top of analyzing gene expression levels per position, quantitative techniques on “pattern” recognition are required to decipher spatial relations between sub/cellular elements. As with our work, multiscale descriptors extracted from structural features in these settings could help elucidate a myriad of underlying mechanisms associated with pattern formation. Our pipeline is well-positioned to facilitate the use of TDA to extract these descriptors from fluorescence microscopy images because of its general-purpose, modular, and accessible design. Moreover, the pipeline possesses general applicability toward the patterns acquired both *in vitro* and *in silico* images, RNA profiling, and medical images.

## Methods

### Cell Culture and Differentiation

The PGP-GATA6 hiPSC line was a gift from Ron Weiss (MIT)^36^; please refer to this source for further biological material availability. Cells were maintained in mTeSR Plus (STEMCELL Technologies) at 37°*C* and 5% CO_2_ with media being changed every other day and cultures being passaged at 80% confluency. Cell culture plates were coated with Matrigel (Growth Factor Reduced, Corning) diluted in KnockOut DMEM (Gibco) for 1*hr* at 37°*C*. During the passage, cells were treated with Accutase (STEMCELL Technologies) for 3*min* at 37°*C*. The dissociated cells were put in PBS and harvested with centrifuging at 200 g for 5*min*. The pellet was resuspended in mTeSR Plus with 10*mM* Y-27632 Dihydrochloride (STEMCELL Technologies) at a final concentration of 10*µM*, and the fraction of resuspension was seeded.

Differentiation experiments were performed at an initial cell seeding density of 25, 000 cells per cm^2^ in mTeSR Plus with 10*µM* Y-27632. The next day, the media was changed to mTeSR Plus with Doxycycline (Sigma-Aldrich) diluted at different concentrations from 0, 5, 15, and 25 ng/ml respectively, and replaced daily for 3 days. The experiments had three replicates per Dox concentration (0, 5, 15, or 25 ng/ml) and antibody choice that detects GATA6 expression (pan-GATA6 or HA).

### Immunofluorescence and Imaging

Cells were grown and treated on Matrigel-coated *µ*-Slide 8 Well chamber slide (ibidi) and fixed for 10*min* in 4% formaldehyde diluted in PBS at room temperature. Each chamber was washed three times in PBS for 5*min*. Cells were then permeabilized for 10*min* in 0.1% Triton X-100 dissolved in PBS with a subsequent washing step. Cells were blocked for 1*hr* in Odyssey buffer (LI-COR Biosciences) and incubated with the primary antibodies overnight at 4°*C*, where each dilution factor varied over antibodies. The next day, each chamber was washed three times in PBS for 5*min*, and the cells were incubated with the secondary antibodies for 30*min* at room temperature. Each chamber was washed three times in PBS for 5*min*. As a final step, cells were stained with Hoechst 33342 (Invitrogen) for 5*min* followed by one wash and the slide was ready for imaging with PBS.

Primary antibodies are Nanog (Cell Signaling Technology 4893S, 1:2000), Gata6 (Abcam ab22600, 1:200, R&D Systems AF1700, which was only used in co-staining, 1:200), and HA (Novus Biologicals NB600-363R, 1:200). Secondary antibodies are donkey anti-mouse Alexa Fluor™Plus 488, donkey anti-rabbit Alexa Fluor™546 (Invitrogen, 1:1000), and donkey anti-goat Alexa Fluor™647 (Invitrogen, 1:1000). Images of 6 *×* 6 multiple fields were acquired by PerkinElmer UltraVIEW VoX Spinning Disk Confocal Microscope and Volocity software (Quorum Technologies) at 10X objective and stitched together with 10% overlap between each tile to generate a large image. To avoid stitching artifacts, we applied brightness and shading corrections when stitching. Images were obtained from five different locations in every replicate. When acquiring the images, the Volocity software converts real-size measurements to pixel values, 1.45*µ*m to 1 pixel at 10X objective. Additionally, before starting the protocol to treat the cells with or without Dox, we obtained the images of the initial condition using Nikon CSU-W1 Spinning Disk Confocal Microscope and the image acquisition software called NIS-Elements. The images were obtained at 10X objective and stitched with 10% overlap between each tile, converting 1.3*µ*m to 1 pixel as real-size measurements to pixel values at the objective. Using Volocity, the stitched large images in the figures were processed with the gamma settings (gamma of 1.5) after brightness enhancement. However, all the quantitative analyses were performed with raw image files.

### Labeling Cells and Signal Intensities

We adapted an existing, histogram-thresholding based cell segmentation pipeline^38^ to quantify cell-specific signal intensities from immunofluorescence microscopy images. A nuclear-localized stain (e.g. Hoechst or DAPI) must be used with this pipeline to identify relative cell locations and to measure other signal intensities. We normalize non-nuclear signals (e.g. Nanog, Gata6, and HA) based on this nuclear-localized stain to account for physical factors (e.g. z-positioning) that affect how signal intensities are detected on a microscope. Our main modifications to the existing segmentation pipeline allow for the consecutive processing of multiple microscopy images across different treatment conditions. This module in our pipeline produces a CSV file for each image which contains cell locations, signal intensities, and other information.

### Signal Thresholding and Cell Type Identification

The cell type identification module of our pipeline generates a signal intensity threshold for each input marker (e.g. HA) based on a user-defined percentile value *n* and a baseline microscopy image set. Each threshold is equal to the *n*th percentile of the corresponding signal intensity distribution from the baseline images. The module uses these thresholds on *all* input images to determine whether each of a cell’s signal intensities should be considered “high/positive” or “low/negative”. Then, this module assigns these discretized values to the cell by updating the CSVs produced in the segmentation module, thereby classifying each cell into a specific cell type. The set of baseline images may be the entire set of input microscopy images or a proper subset.

In our work, we generated thresholds for NANOG, pan-GATA6, and HA expression while considering that signal intensities may be affected by the choice of imaging location in each well replicate. We partitioned our microscopy images into subsets according to the red marker (pan-GATA6 or HA) and imaging location and then generated green (NANOG) and red thresholds for each subset using the same percentile value *n*. For each subset, we chose the images from the highest Dox treatment group (25 ng/ml) as our baseline image set for the green threshold given that NANOG expression is expected to be lowest when GATA6 expression is highest. We chose the images from the lowest Dox treatment group (0 ng/ml) as our baseline image set for the red thresholds for the opposite reason.

### Percentile Threshold and Patch Size Selection

We discuss the following parameters chosen for our microscopy images: percentile threshold and patch size. Signal intensity thresholds rely on a user-defined percentile *n*, so the choice of *n* affects the population distributions of each cell type. To determine which *n* to use, we considered how the population distributions of the four cell types change when *n* = 65, 70, 75, 80, 85 using the cell type identification module of our pipeline. We report this information in **Supplementary Table** S1. Our goal in choosing *n* was to maximize the number of cells classified as R^*−*^G^+^ or R^+^G^*−*^ across both red markers (pan-GATA6 and HA) because NANOG and GATA6 are indicators for pluripotency and differentiation, respectively. The *n* that maximized this number was directly proportional to the Dox concentration in general, so we chose middle value *n* = 75 for all Dox concentrations.

Our choice of patch size balances (i) the presence of big, rare regions devoid of cells and (ii) the ratio of empty patches for each cell type. Large patches may anomalous, vacant regions which might not be representative of the multicellular patterning associated with differentiation; see **Supplementary Fig**. S13 for example. On the other hand, splitting images into smaller patches increases the number of empty patches in every cell type (**Supplementary Table** S7). For instance, the number of empty patches of R^*−*^G^+^ cell type in the pan-GATA6 group is 43.3% when there every image is partitioned into 25 patches.

### PLs for Patches of Discretized Microscopy Images

Each discretized, 2, 850 *×* 2, 850 pixel (4, 132.5 *×* 4, 132.5 *µ*m) microscopy image was split into 16 even non-overlapping square patches. For each patch, we considered each cell type individually and applied the following TDA pipeline. We computed the Delaunay filtration, persistence diagrams, and representative cycles using R package ‘TDA’^69^, which wraps C++ libraries ‘GUDHI’^70^ and ‘Dionysus’^71^. PL computations used R package ‘tda-tools’^72^, which wraps the persistence landscape toolbox^73^.

PLs are defined as follows. For *b < d*, consider the function *f*_*b,d*_ : ℝ *→* ℝ defined by *f*_*b,d*_(*t*) = *t −b* if *b* ≤ *t* ≤ (*b* + *d*)*/*2; *f*_*b,d*_(*t*) = *d −t*, if (*b* + *d*)*/*2 *≤ t ≤ d*; and *f*_*b,d*_(*t*) = 0 otherwise. Let *𝒟* be a persistence diagram. Its corresponding PL is given by the sequence of functions (*λ*_*k*_), *k* = 1, 2, 3, …, where *λ*_*k*_ : ℝ *→* ℝ is given by defining *λ*_*k*_(*t*) to be the *k*th-largest value of *f*_*b,d*_(*t*) over the points (*b, d*) in *𝒟* . The function *λ*_*k*_ is called the *k*th PL function of *𝒟* and the parameter *k* is referred to as its depth. We visualize the PL by graphing the PL functions on the same plot and depicting them with a set of colors that repeat at every 15 depths.

We discretized the PLs using a grid with a step size equal to 0.3, which is fine enough to capture fine spatial features and provide nice visualizations. We use all depths of the PLs that are nonzero for plots. However, we used PLs up to depth 30 which we concatenated to obtain a single vector for computations. As a result, we obtained four vectors for every patch, one for each cell type.

### Statistics

To check for a statistically significant difference between two groups, we used the permutation test (one-tailed)^74^ on PL vectors. Each of the Dox treatment groups consisted of 240 PL vectors, while the group consisting of images taken at the start of our treatment protocol had only 48 PL vectors. A permutation test is a non-parametric test on the two or more samples which is often appropriate when the underlying distributions of the samples are unknown^74^. After choosing a test statistic for the permutation test, the *observed* test statistic is computed using the samples as given. Then, the members from both samples are combined and assigned to the two groups randomly, called a *shuffle*. This is repeated *n* times and the test statistic is recomputed after each shuffle. Our test statistic was the Euclidean distance between the means of the PL vectors of the two groups, and we shuffled the PL vectors 10, 000 times. Under the assumption that both groups cannot be distinguished by the test statistic, the p-value was estimated as the proportion of permutations for which the distance was at least as large as the observed distance.

We used the threshold of 5E*−*2 to determine the significance of the p-value.

For comparison, we also performed k-means clustering and comparisons of local density using DBSCAN^46^. DBSCAN uses the neighborhood density of each data point to identify clusters and is robust to noise and outliers. To compute the local density, we used the positional information (i.e., *X* and *Y* coordinates) of each cell in each group. We selected the minimum number of points (*MinPts* = 5) for identifying a dense region and then employed the kneedle algorithm^75^ to determine the optimal radius (*ε*) of the neighborhood around a data point. This algorithm detects a knee in the curve plotting the distance of all points to the corresponding *k*-nearest neighbor (*k* = *MinPts−* 1) per each image. In addition to calculating all local densities, we obtained a mean local density for every image and compared each instance between groups using Welch’s two-sample t-test. Statistical differences were also determined using a significance level of 5E*−*2. K-means clustering is an unsupervised algorithm that partitions all data points into *k* number of clusters to classify and identify cohorts in a given dataset. For every image, we computed the means of normalized NANOG and pan-GATA6 or HA fluorescent intensities of certain cell types and assigned these to each data point along with the corresponding mean local density. After consolidating all the data points (i.e. means of intensity profiles and local densities per image) for each group, we used the gap static method to determine the optimal number of clusters.

### Machine Learning

For our machine learning computations, we concatenated the PL vectors of the four cell types to obtain a single vector of length 127, 770 for each patch. For every PL vector, we removed the coordinates that were zero. Next, we normalized these vectors coordinate-wise; for each coordinate, we subtracted the mean and divided it by the standard deviation. Afterward, we used normalized PLs as inputs for machine learning.

For binary classification, we used support vector machines (SVM). When there were more than two classes, we used multiclass SVM using the “one-against-one” approach^76^. In this method, binary classifiers are trained for every pair of classes. The predicted class is the one that was chosen the most often by these classifiers. We performed our computations using the R package ‘kernlab’^77^, using default settings and cost equal to 10. To estimate SVM model accuracy, we used 5-fold cross-validation in the binary case and 10-fold cross-validation in the multiclass case. In each case, we repeated the classification 20 times and reported the average accuracy.

For regression, we used a variant of SVM called support vector regression (SVR)^76^ using the *ε*-insensitive loss function. For this loss function, the value of *ε* defines a margin of tolerance where no penalty is given to errors. We chose *ε* = 0.01 and otherwise used the same hyper-parameters as with SVM. We also used 10-fold cross-validation with 20 repetitions and plotted the average predicted values as box plots (**Fig**. 4 and **Supplementary Fig**. S5). A box plot provides information about the distribution of data based on five quartiles (excluding outliers): 100th (tip of top line), 75th (top of box), 50th (horizontal line), 25th (bottom of box), and 0th (tip of bottom line). The interquartile range is the difference between the 75th and 25th percentiles. Gray points in a box plot are outliers.

## Data Availability

The microscopy images associated with our work, including an example image set, are publicly available on Figshare: https://figshare.com/projects/TDA_Microscopy_Data/148855.

## Code Availability

Our computational pipeline is freely available via the Apache 2.0 license on GitHub along with example scripts: https://github.com/kemplab/TDA-Microscopy-Pipeline.

## Acknowledgements

The authors would like to thank Hector Baños, Alex Elchesen, Afaf Saidi, and Ashleigh Thomas for helpful discussions at the initial stage of this project. The authors also thank the members of the Laboratory for Systems Medicine at the University of Florida for their feedback during manuscript revisions. This research was supported by the NSF-Simons Southeast Center for Mathematics and Biology (SCMB) through the grants National Science Foundation DMS1764406 and Simons Foundation/SFARI 594594.

## Author contributions statement

All authors collaborated in designing research. E.P. conducted the experiments and imaging. E.P and J.T. preprocessed the experimental data. I.H. and E.P. performed quantitative data analysis. I.H., E.P., and D.A.C. discussed and interpreted the results. I.H. and J.T. implemented the computational pipeline; P.B. and D.A.C. contributed code and commentary. All authors discussed the results and wrote the manuscript.

## Additional information

**Competing interests** The authors declare no competing interest.

