## Supplementary Information for "Topological data analysis of pattern formation of human induced pluripotent stem cell colonies"

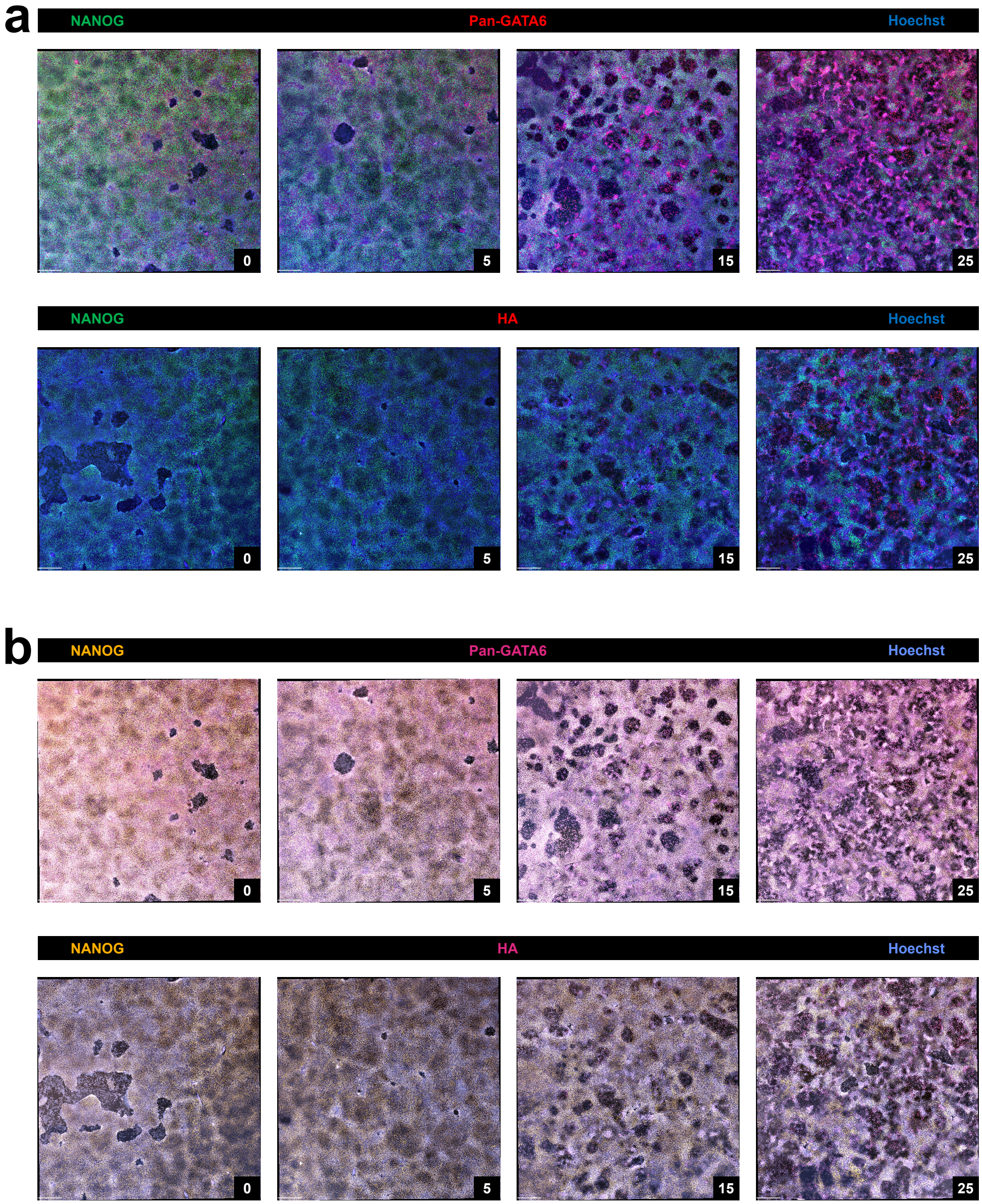


Fig. S1: Spatial organization of the genetically engineered hiPSC line with various Dox concentrations. **a** Immunofluorescence images of NANOG and pan-GATA6 or HA at different Dox concentrations of 0, 5, 15, and 25 ng/ml (scale bar, 440$\mu m$). The treatments with higher Dox concentrations induce more GATA6-HA expression. **b** The images are converted to a color palette accessible for color vision deficiency [1]. Using Volocity, we applied the gamma changes (gamma of 1.5) after brightness enhancement on all stitched large images used in the figure to have better contrast for representation.

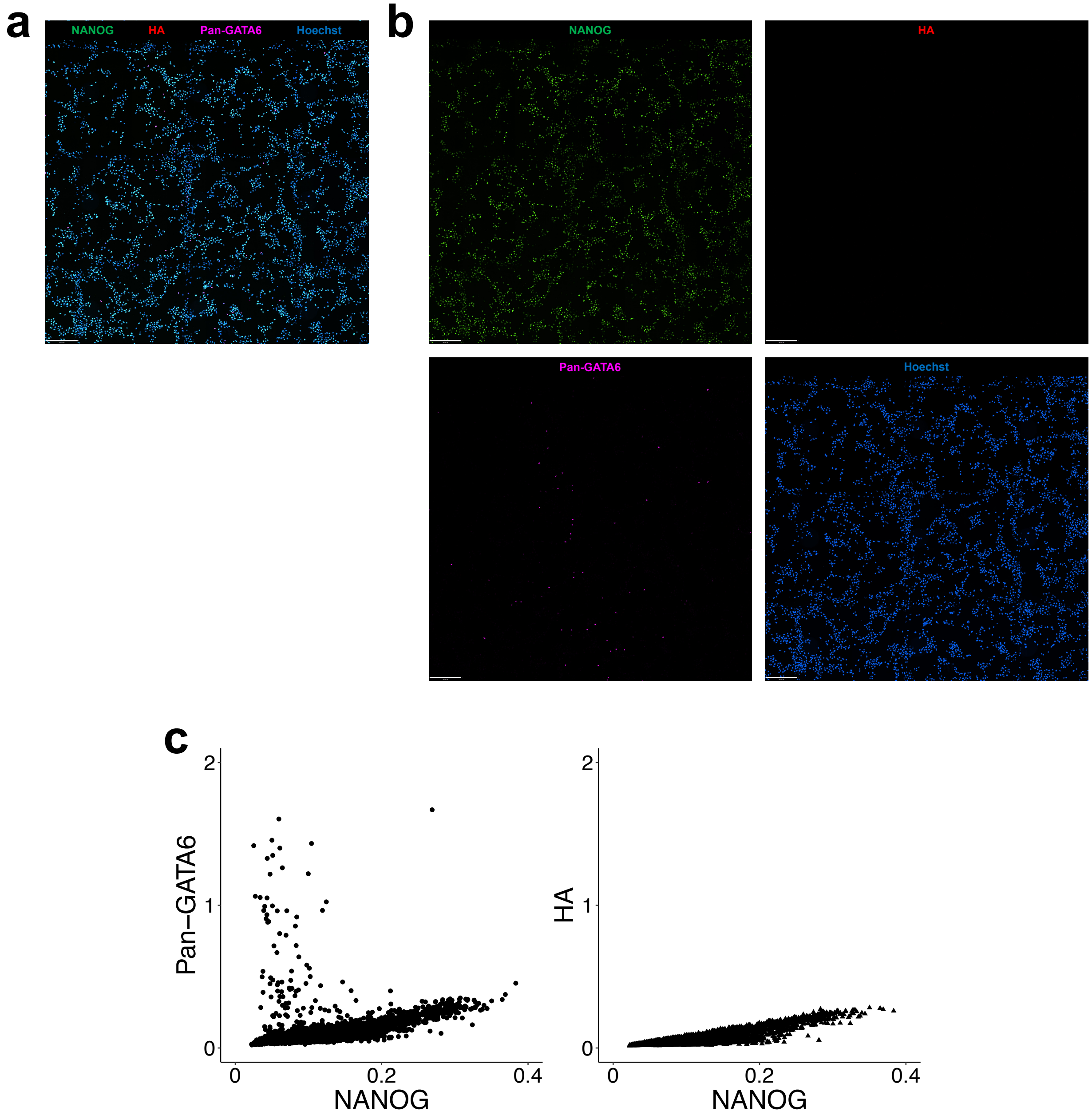


Fig. S2: Initial pattern formation before any Dox treatments. **a** Immunofluorescence image of NANOG, HA, and pan-GATA6 (scale bar, 390$\mu m$). **b** Separated images of single channels to show each fluorescent marker respectively. Using Volocity, we applied the gamma changes (gamma of 1.5) after brightness enhancement on the stitched large image used in this figure to have better contrast for representation. **c** Quantification of the segmented image by each channel. Fluorescent intensities of NANOG, HA, and pan-GATA6 are normalized to the corresponding nuclear Hoechst value.

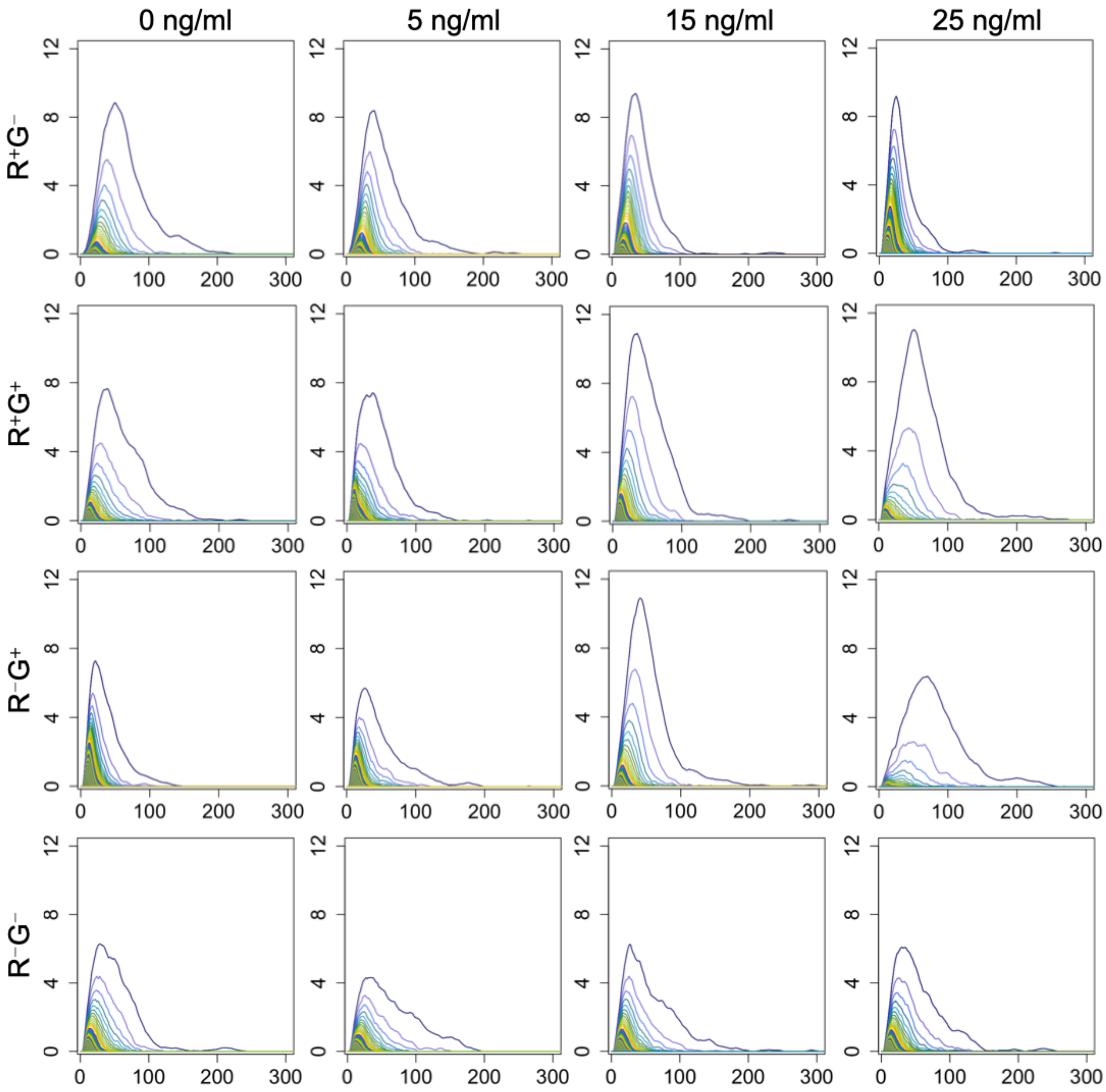


Fig. S3: Comparison of the average persistence landscapes across various Dox concentrations for all cell types in the pan-GATA6 group. As Dox concentration increases the average persistence landscape is getting taller and narrower for $R^+G^-$ cell type. Different trends of the average persistence landscape's behavior are witnessed for other cell types.

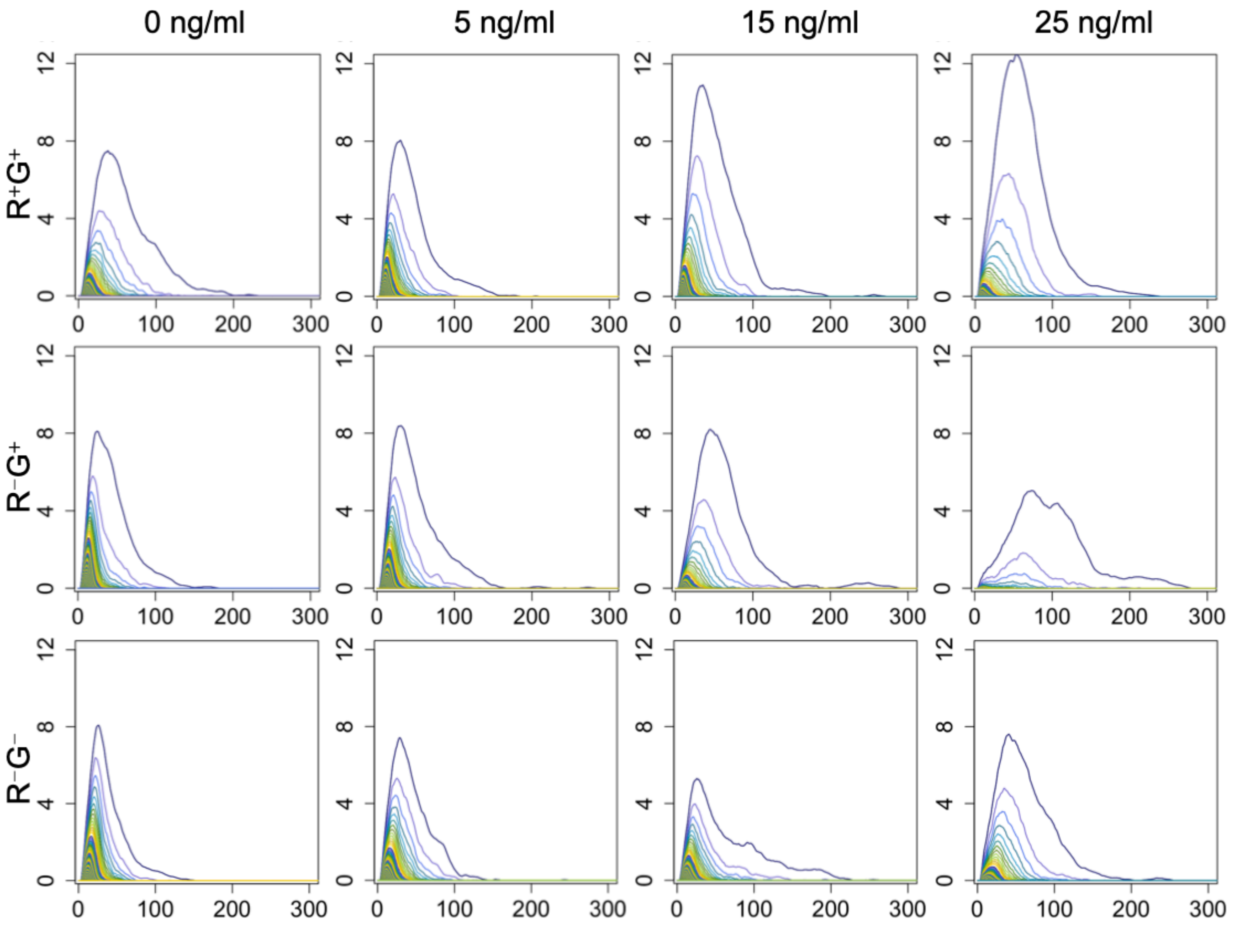


Fig. S4: Comparison of the average persistence landscapes across various Dox concentrations for $R^+G^+$, $R^-G^+$, $R^-G^-$ cell types in the HA group. As Dox concentration increases the average persistence landscape is getting shorter and wider for $R^-G^+$ cell type. Different trends of the average persistence landscape's behavior are witnessed for other cell types.

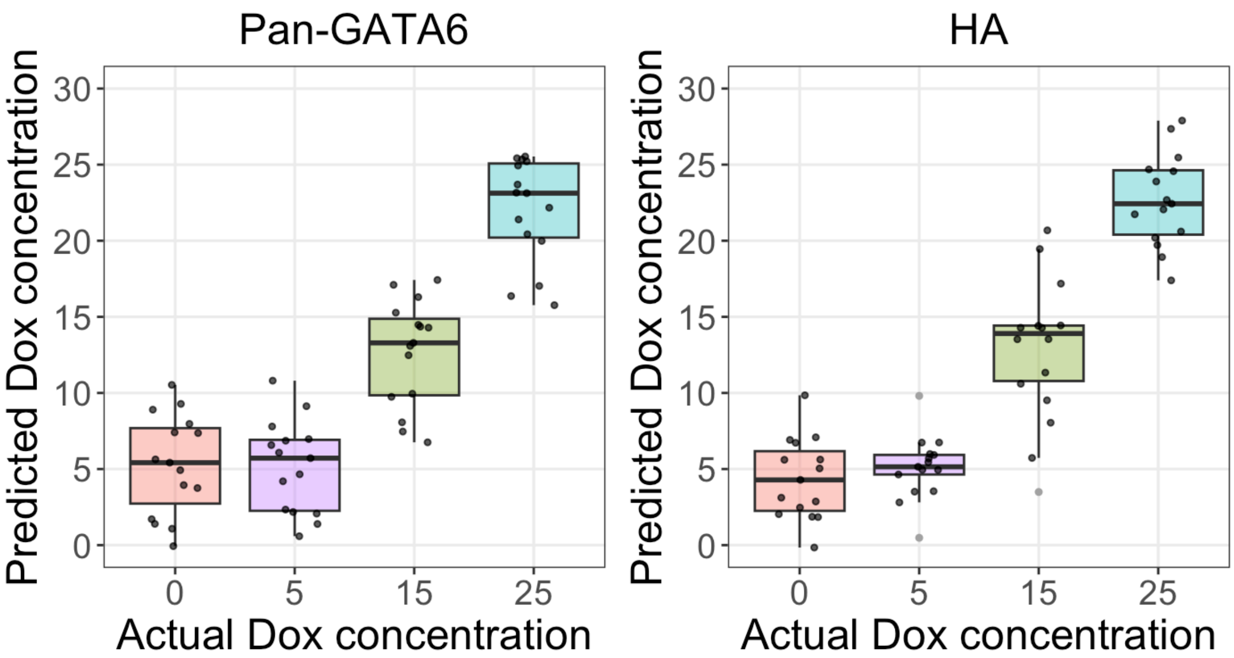


Fig. S5: Support vector regression was performed on the cell count vectors extracted from patches, and then, for every image, the Dox concentration predictions of its patches were averaged. The image predictions of each Dox treatment group are horizontally spread apart for better visualization, and the outliers are marked with gray points. Overall the image predictions are far from their actual Dox concentration.

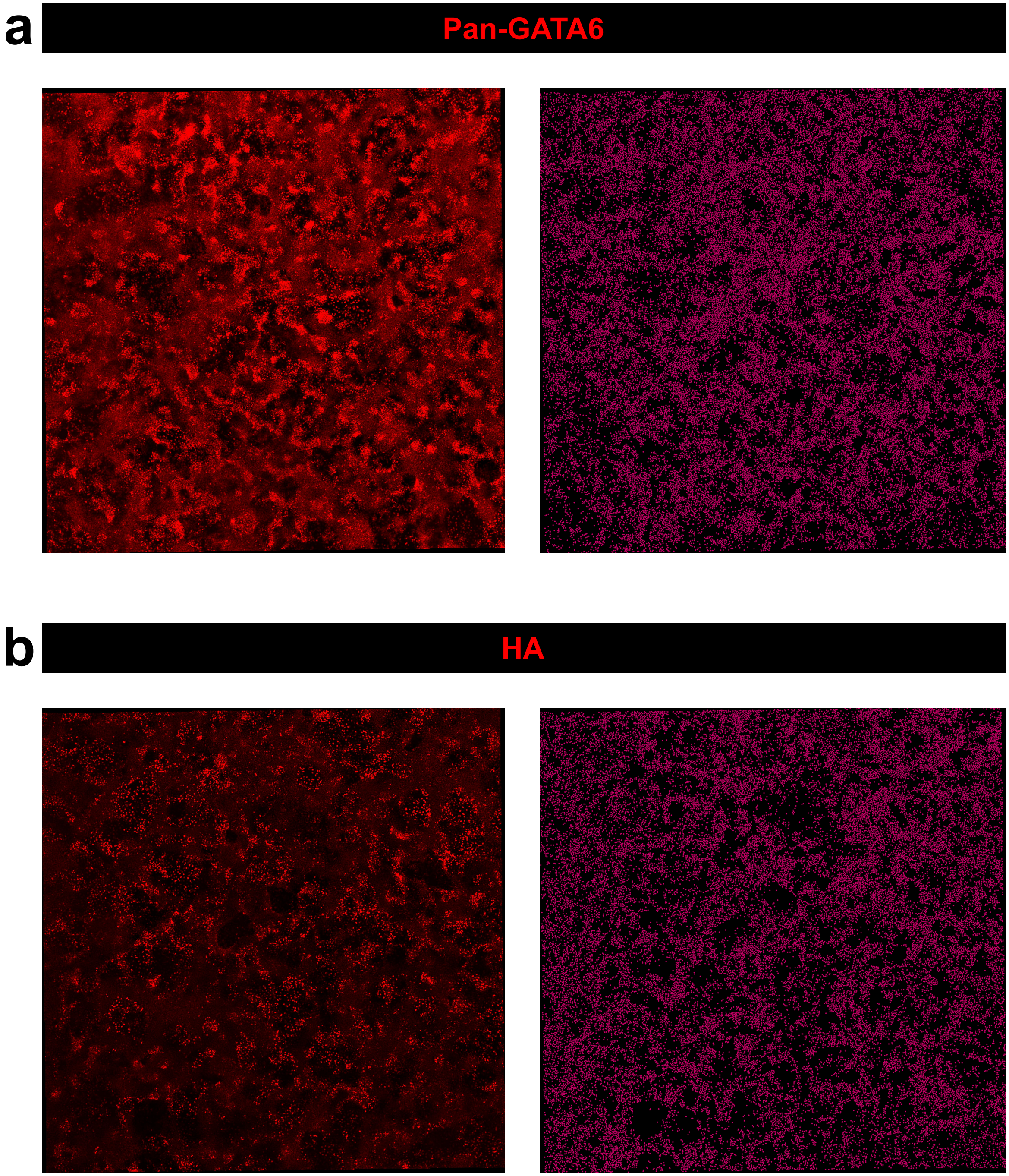


Fig. S6: Different patterns between pan-GATA6 and HA populations at 25 ng/ml Dox concentration. At each panel of **a** and **b**, the confocal microscopy images (left) show the patterns with the channel of pan-GATA6 or HA only present. The right side of the panels displays its corresponding discretized image of the $R^+G^-$ cell type. In general, the HA group **b** has a larger number and size of empty regions than the pan-GATA6 **a**. Using Volocity, we applied the gamma changes (gamma of 1.5) after brightness enhancement on all stitched large images used in the figure to have better contrast for representation.

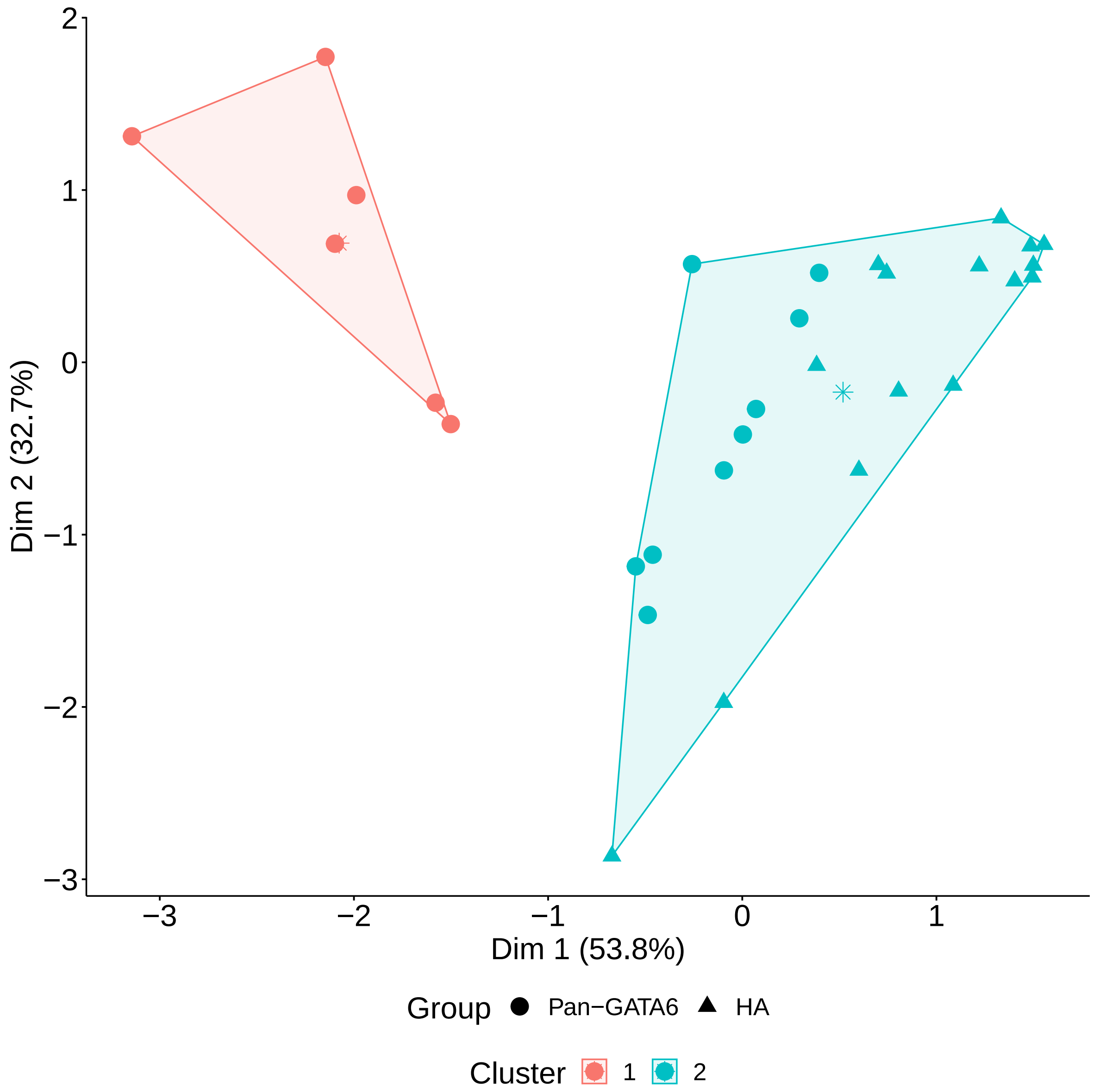


Fig. S7: K-means clustering results between pan-GATA6 and HA populations, using the $R^+G^-$ cells in the 25 ng/ml Dox treatment groups. We used the gap statistic method for computing the optimal number of clusters, and an asterisk on each cluster is its centroid. The clustering results do not show clear partitions between pan-GATA6 and HA groups since data points from each group are aggregated in the same cluster.

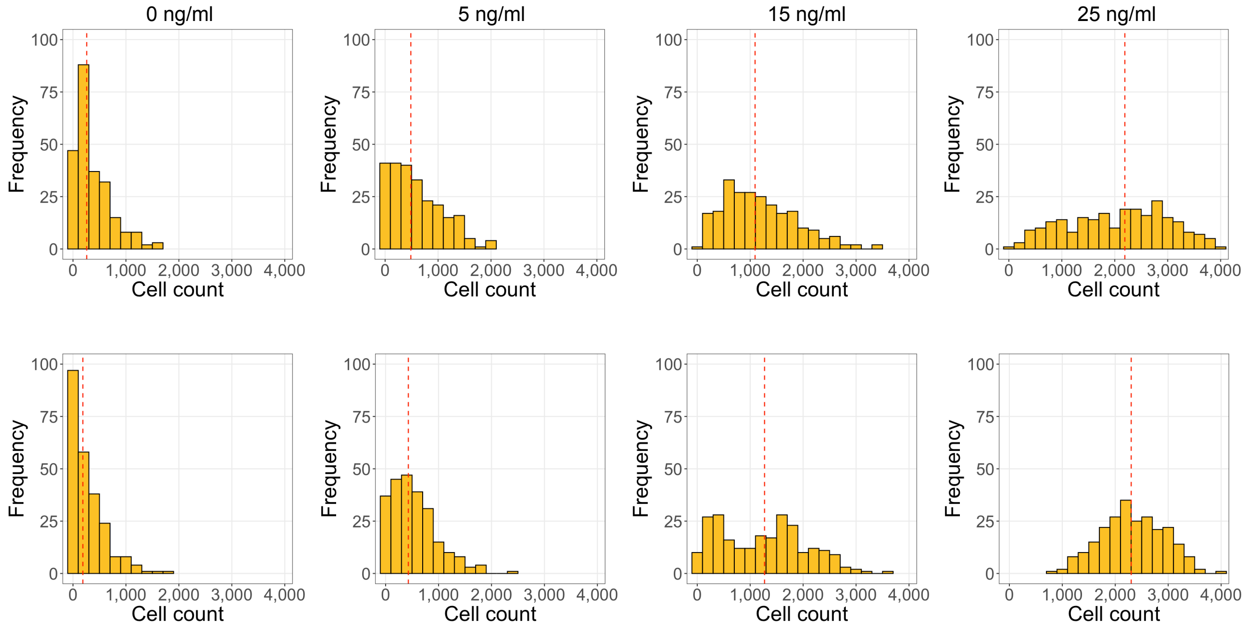


Fig. S8: Histograms for each Dox treatment group of the cell counts per patch for $R^+G^-$ cell type in pan-GATA6 (top row) and HA (bottom row) populations. Red dashed lines indicate medians. The cell counts increase as Dox concentration gets higher.

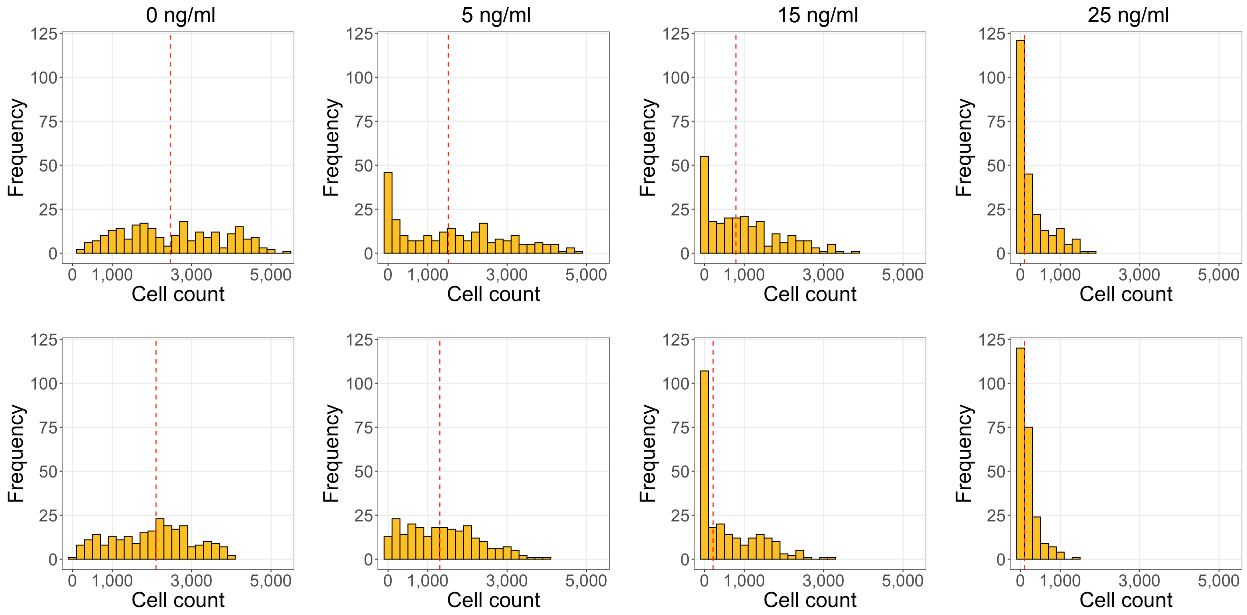


Fig. S9: Histograms for each Dox treatment group of the cell counts per patch for $R^{-}G^{+}$ cell type in pan-GATA6 (top row) and HA (bottom row) populations. Red dashed lines indicate medians. The cell counts decrease as Dox concentration gets higher.

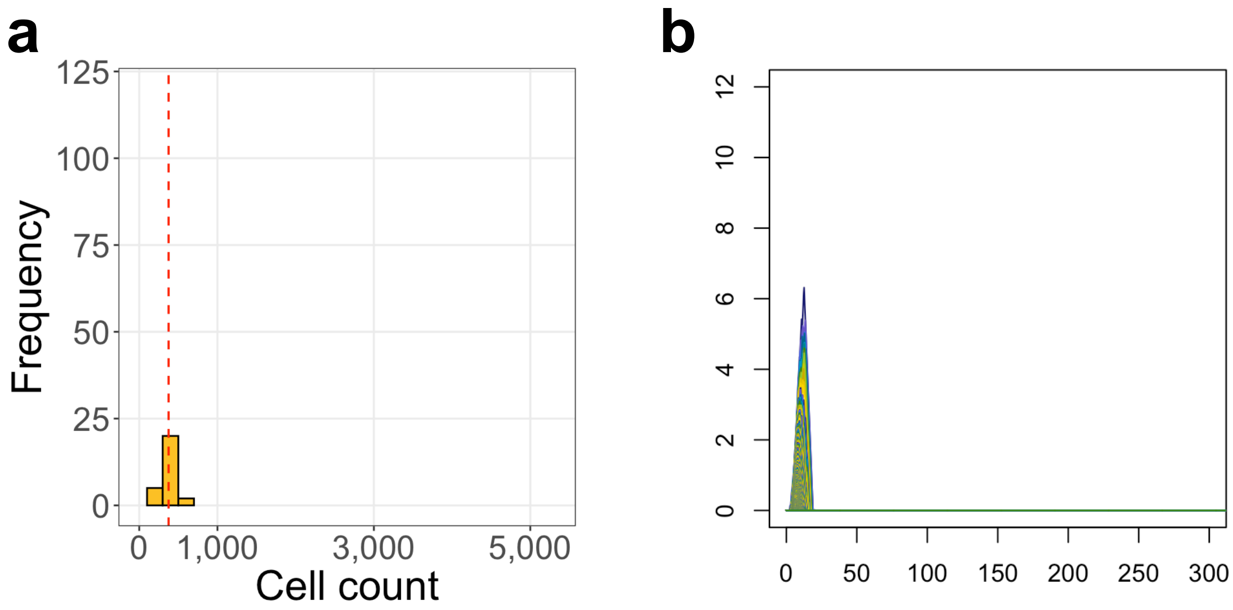


Fig. S10: **a** Histogram of the cell counts in the initial condition group. Red dashed line indicates the median. **b** Average persistence landscape of the initial condition group.

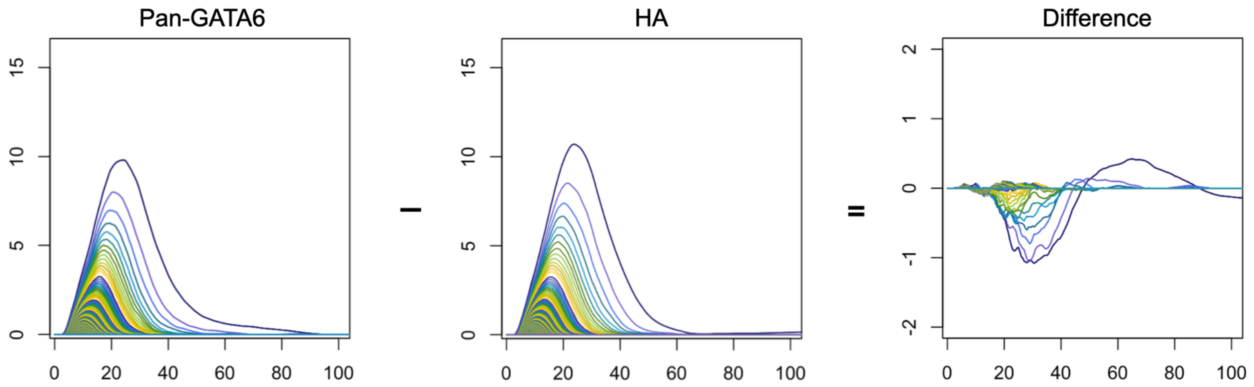


Fig. S11: Average persistence landscapes of the pan-GATA6 and HA populations for $R^+G^-$ cell type based on patches with more than $1,000$ cells and their difference. The negative region in the difference plot indicates that there are more persistence holes in HA group than in pan-GATA6 group.

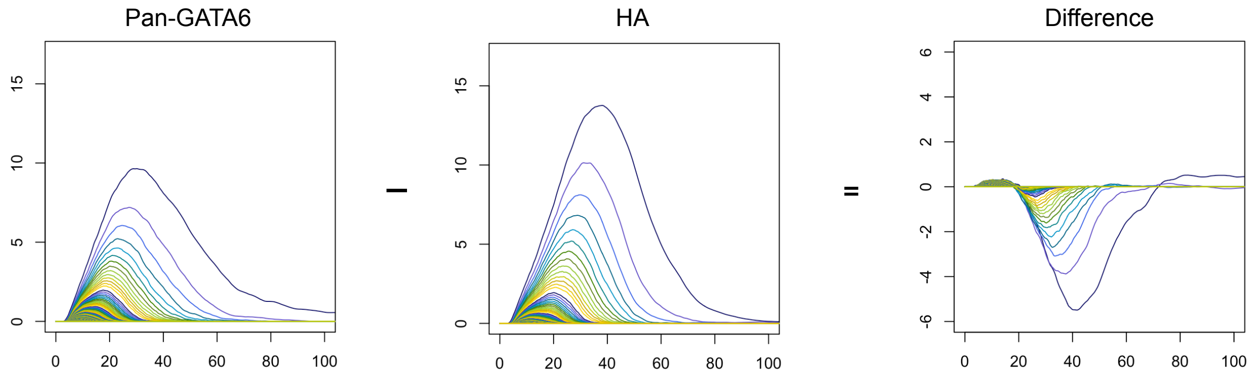


Fig. S12: Average persistence landscapes of the pan-GATA6 and HA populations for $R^{+}G^{-}$ cell type using stitched image files without brightness and shading corrections (see *Methods*). The negative region in the difference plot indicates that there are more persistence holes in HA group than in pan-GATA6 group.

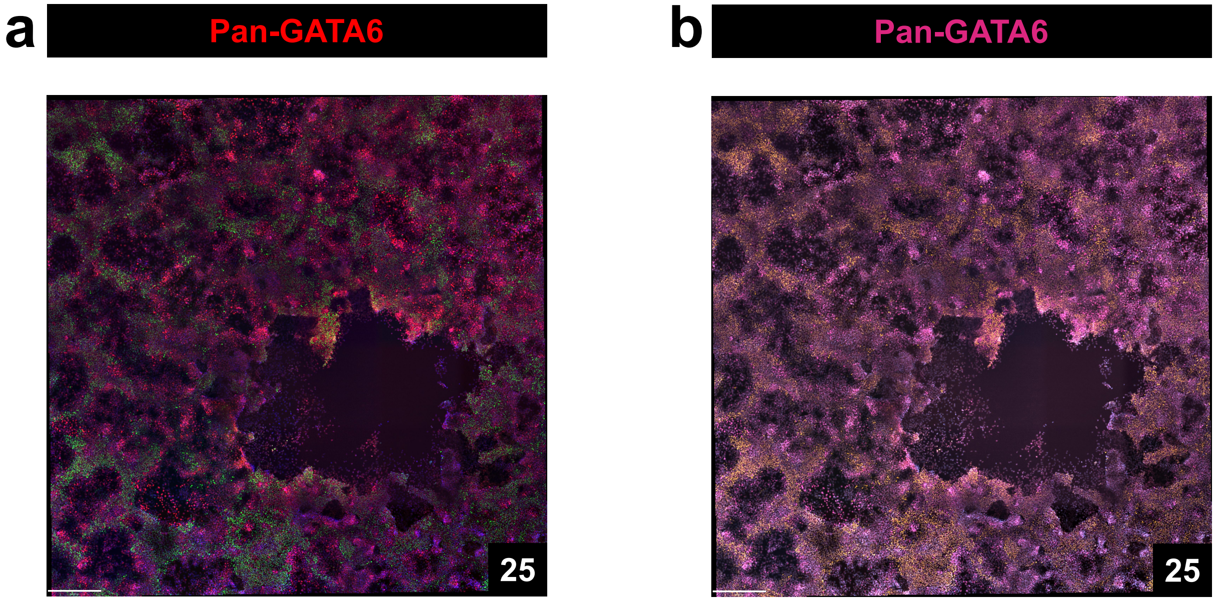


Fig. S13: An example of a confocal microscopy image that has a large region devoid of cells. **a** Immunofluorescent image of NANOG and pan-GATA6 at 25 ng/ml Dox concentration (scale bar, 440$\mu m$). Dividing the image into large patches can include the majority of the empty area, which may not represent the differentiation patterns of cell colonies. **b** Converted image with the same color palette used in **Fig.** S1 [1]. Using Volocity, we applied the gamma changes (gamma of 1.5) after brightness enhancement on all stitched large images used in the figure to have better contrast for representation.

Table S1: Percentage distributions of the four cell types ($R^-G^-$, $R^-G^+$, $R^+G^-$, and $R^+G^+$) for all pan-GATA6 and HA populations, organized with respect to marker, percentile threshold, and Dox concentration. Percentages are rounded to the nearest two decimal points. For each combination of marker and Dox concentration, we also report the total number of cells across all well replicates and imaging locations.

| Marker | Percentile | Dox | Cell Total | $R^-G^-$ | $R^-G^+$ | $R^+G^-$ | $R^+G^+$ |
|---|---|---|---|---|---|---|---|
| Pan-GATA6 | 65$^{th}$ | 0 ng/ml | 1,312,524 | 15.31% | 48.49% | 5.71% | 30.49% |
| | | 5 ng/ml | 1,357,299 | 11.07% | 28.25% | 7.61% | 53.07% |
| | | 15 ng/ml | 1,219,235 | 15.40% | 20.22% | 18.30% | 46.09% |
| | | 25 ng/ml | 1,071,314 | 21.50% | 7.54% | 42.70% | 28.25% |
| | 70$^{th}$ | 0 ng/ml | 1,312,524 | 20.21% | 48.05% | 6.32% | 25.41% |
| | | 5 ng/ml | 1,357,299 | 14.37% | 28.84% | 8.90% | 47.88% |
| | | 15 ng/ml | 1,219,235 | 19.08% | 19.98% | 20.69% | 40.25% |
| | | 25 ng/ml | 1,071,314 | 24.65% | 7.17% | 44.74% | 23.45% |
| | 75$^{th}$ | 0 ng/ml | 1,312,524 | 26.69% | 46.11% | 6.90% | 20.30% |
| | | 5 ng/ml | 1,357,299 | 18.67% | 28.58% | 10.50% | 42.25% |
| | | 15 ng/ml | 1,219,235 | 23.69% | 19.08% | 23.21% | 34.02% |
| | | 25 ng/ml | 1,071,314 | 28.30% | 6.60% | 46.14% | 18.96% |
| | 80$^{th}$ | 0 ng/ml | 1,312,524 | 35.17% | 42.21% | 7.44% | 15.18% |
| | | 5 ng/ml | 1,357,299 | 24.32% | 27.23% | 12.58% | 35.87% |
| | | 15 ng/ml | 1,219,235 | 29.46% | 17.50% | 25.55% | 27.49% |
| | | 25 ng/ml | 1,071,314 | 32.56% | 5.88% | 46.83% | 14.73% |
| | 85$^{th}$ | 0 ng/ml | 1,312,524 | 45.87% | 36.27% | 7.71% | 10.14% |
| | | 5 ng/ml | 1,357,299 | 31.75% | 24.63% | 15.12% | 28.50% |
| | | 15 ng/ml | 1,219,235 | 36.57% | 15.32% | 27.39% | 20.72% |
| | | 25 ng/ml | 1,071,314 | 37.89% | 5.01% | 46.36% | 10.74% |
| HA | 65$^{th}$ | ng/ml 0 | 1,235,476 | 21.56% | 42.47% | 4.53% | 31.43% |
| | | 5 ng/ml | 1,296,037 | 18.76% | 26.71% | 7.71% | 46.82% |
| | | 15 ng/ml | 1,152,243 | 18.02% | 13.98% | 20.83% | 47.17% |
| | | 25 ng/ml | 1,012,962 | 13.12% | 4.57% | 49.39% | 32.92% |
| | 70$^{th}$ | 0 ng/ml | 1,235,476 | 26.65% | 41.44% | 4.96% | 26.96% |
| | | 5 ng/ml | 1,296,037 | 23.15% | 26.59% | 8.66% | 41.60% |
| | | 15 ng/ml | 1,152,243 | 21.14% | 13.43% | 23.14% | 42.29% |
| | | 25 ng/ml | 1,012,962 | 15.38% | 4.51% | 52.52% | 27.58% |
| | 75$^{th}$ | 0 ng/ml | 1,235,476 | 33.46% | 38.72% | 5.50% | 22.32% |
| | | 5 ng/ml | 1,296,037 | 29.00% | 25.41% | 9.84% | 35.76% |
| | | 15 ng/ml | 1,152,243 | 24.98% | 12.35% | 25.73% | 36.94% |
| | | 25 ng/ml | 1,012,962 | 18.13% | 4.29% | 55.39% | 22.19% |
| | 80$^{th}$ | 0 ng/ml | 1,235,476 | 42.30% | 34.12% | 6.02% | 17.56% |
| | | 5 ng/ml | 1,296,037 | 36.75% | 22.90% | 11.05% | 29.30% |
| | | 15 ng/ml | 1,152,243 | 29.70% | 10.73% | 28.41% | 31.16% |
| | | 25 ng/ml | 1,012,962 | 21.52% | 3.91% | 57.48% | 17.09% |
| | 85$^{th}$ | 0 ng/ml | 1,235,476 | 47.52% | 30.59% | 5.47% | 16.43% |
| | | 5 ng/ml | 1,296,037 | 40.09% | 23.50% | 11.95% | 24.46% |
| | | 15 ng/ml | 1,152,243 | 31.64% | 10.53% | 29.24% | 28.59% |
| | | 25 ng/ml | 1,012,962 | 22.37% | 3.95% | 57.82% | 15.86% |

Table S2: Permutation test results on the pairwise Euclidean distances between persistence landscape vectors of initial condition and Dox treatment groups. The hypothesis testing was performed comparing all cells in the initial condition group against $R^-G^+$ cells in both the pan-GATA6 and HA groups. The p-values were computed using 10,000 permutations.

| Marker | Dox concentration | Initial condition |
|---|---|---|
| Pan-GATA6 ($R^-G^+$ cells) | 0 ng/ml | 1E-4 |
| | 5 ng/ml | 1E-4 |
| | 15 ng/ml | 1E-4 |
| | 25 ng/ml | 1E-4 |
| HA ($R^-G^+$ cells) | 0 ng/ml | 1E-4 |
| | 5 ng/ml | 1E-4 |
| | 15 ng/ml | 1E-4 |
| | 25 ng/ml | 1E-4 |

Table S3: Permutation test results on the pairwise Euclidean distances between persistence landscape vectors of two distinct Dox treatment groups. The hypothesis testing was performed separately for each cell type in both the pan-GATA6 and HA groups. The p-values were computed using $10,000$ permutations.

| Marker | Cell type | 0 vs 5 ng/ml | 0 vs 15 ng/ml | 0 vs 25 ng/ml | 5 vs 15 ng/ml | 5 vs 25 ng/ml | 15 vs 25 ng/ml |
|---|---|---|---|---|---|---|---|
| Pan-GATA6 | $R^+G^-$ | 1E-4 | 1E-4 | 1E-4 | 1E-4 | 1E-4 | 1E-4 |
| | $R^+G^+$ | 1E-4 | 1E-4 | 1E-4 | 1E-4 | 1E-4 | 1E-4 |
| | $R^-G^+$ | 1E-4 | 1E-4 | 1E-4 | 1E-4 | 1E-4 | 1E-4 |
| | $R^-G^-$ | 1E-4 | 9.1E-2 | 3.16E-1 | 1.04E-2 | 2E-3 | 6.424E-1 |
| HA | $R^+G^-$ | 1E-4 | 1E-4 | 1E-4 | 1E-4 | 1E-4 | 1E-4 |
| | $R^+G^+$ | 1E-4 | 1E-4 | 1E-4 | 1E-4 | 1E-4 | 1E-4 |
| | $R^-G^+$ | 2E-4 | 1E-4 | 1E-4 | 1E-4 | 1E-4 | 1E-4 |
| | $R^-G^-$ | 1E-4 | 1E-4 | 1E-4 | 1E-4 | 1E-4 | 1E-4 |

Table S4: Confusion matrices of a single instance of multiclass support vector machines with 10-fold cross validation on the persistence landscape vectors and the cell count vectors of different Dox treatment groups in the pan-GATA6 and HA populations. The yellow/orange/red color scheme corresponds to low/medium/high numbers, respectively.

| | Marker | Actual Dox concentration | Predicted Dox concentration | | | |
|---|---|---|---|---|---|---|
| | | | 0 ng/ml | 5 ng/ml | 15 ng/ml | 25 ng/ml |
| PLs | Pan-GATA6 | 0 ng/ml | 136 | 91 | 20 | 1 |
| | | 5 ng/ml | 89 | 136 | 12 | 1 |
| | | 15 ng/ml | 14 | 12 | 192 | 19 |
| | | 25 ng/ml | 1 | 1 | 16 | 219 |
| | HA | 0 ng/ml | 152 | 86 | 20 | 1 |
| | | 5 ng/ml | 72 | 142 | 13 | 0 |
| | | 15 ng/ml | 16 | 12 | 190 | 18 |
| | | 25 ng/ml | 0 | 0 | 17 | 221 |
| Cell counts | Pan-GATA6 | 0 ng/ml | 176 | 108 | 41 | 4 |
| | | 5 ng/ml | 40 | 104 | 23 | 0 |
| | | 15 ng/ml | 21 | 24 | 152 | 22 |
| | | 25 ng/ml | 3 | 4 | 24 | 214 |
| | HA | 0 ng/ml | 138 | 64 | 21 | 0 |
| | | 5 ng/ml | 74 | 154 | 39 | 0 |
| | | 15 ng/ml | 28 | 22 | 147 | 24 |
| | | 25 ng/ml | 0 | 0 | 22 | 216 |

Table S5: Additional information on support vector regression box plots on the persistence landscape vectors and the cell count vectors in the pan-GATA6 and HA groups. Each table contains the lower quartile (Q1), the upper quartile (Q3), median, interquartile range (IQR=Q3-Q1), and the deviation of the median from the actual Dox concentration (error).

| | Marker | | 0 ng/ml | 5 ng/ml | 15 ng/ml | 25 ng/ml |
|---|---|---|---|---|---|---|
| PLs | Pan-GATA6 | Q1 | 2.45 | 1.86 | 12.93 | 22.01 |
| | | Median | 2.84 | 4.32 | 15.68 | 22.55 |
| | | Q3 | 4.37 | 6.38 | 16.45 | 23.89 |
| | | IQR | 1.92 | 4.52 | 3.52 | 1.88 |
| | | Error | 2.84 | 0.68 | 0.68 | 2.45 |
| | HA | Q1 | 1.95 | 2.95 | 11.00 | 22.55 |
| | | Median | 3.57 | 3.79 | 14.64 | 23.24 |
| | | Q3 | 4.10 | 5.36 | 15.40 | 24.38 |
| | | IQR | 3.15 | 2.41 | 4.40 | 1.83 |
| | | Error | 3.57 | 1.21 | 0.36 | 1.76 |
| Cell counts | Pan-GATA6 | Q1 | 2.72 | 2.27 | 9.85 | 20.20 |
| | | Median | 5.42 | 5.71 | 13.31 | 23.13 |
| | | Q3 | 7.70 | 6.93 | 14.88 | 25.08 |
| | | IQR | 4.98 | 4.66 | 5.03 | 4.88 |
| | | Error | 5.42 | 0.71 | 1.69 | 1.87 |
| | HA | Q1 | 2.25 | 4.11 | 10.09 | 20.41 |
| | | Median | 4.28 | 5.13 | 13.53 | 22.44 |
| | | Q3 | 6.19 | 5.96 | 14.41 | 24.65 |
| | | IQR | 3.94 | 1.85 | 4.32 | 4.24 |
| | | Error | 4.28 | 0.13 | 1.47 | 2.56 |

Table S6: Accuracies of pairwise support vector machines with 5-fold cross validation on the persistence landscape vectors and the cell count vectors of two distinct Dox treatment groups in the pan-GATA6 and HA populations. The yellow/orange/red color scheme corresponds to low/medium/high accuracies, respectively.

| | Marker | 0 vs 5 ng/ml | 0 vs 15 ng/ml | 0 vs 25 ng/ml | 5 vs 15 ng/ml | 5 vs 25 ng/ml | 15 vs 25 ng/ml |
|---|---|---|---|---|---|---|---|
| PLs | Pan-GATA6 | 61.81% | 90.52% | 98.66% | 89.55% | 98.64% | 91.85% |
| | HA | 62.59% | 89.83% | 99.65% | 90.93% | 99.71% | 92.06% |
| Cell counts | Pan-GATA6 | 63.61% | 81.61% | 97.00% | 82.36% | 98.23% | 88.48% |
| | HA | 68.19% | 81.83% | 98.95% | 83.69% | 99.11% | 88.14% |

Table S7: The ratio of empty patches in each cell type in both pan-GATA6 and HA groups. The ratio was computed when images were split into 9, 16, and 25 square patches. Note that the larger the number of patches the smaller their size. For a fixed cell type, the ratio of empty patches increases as images get split into smaller patches.

| # of patches | Marker | $R^+G^-$ | $R^+G^+$ | $R^-G^+$ | $R^-G^-$ |
|---|---|---|---|---|---|
| 9 | Pan-GATA6 | 0.8% | 0.4% | 11.7% | 3.3 % |
| | HA | 0.4% | 0.4% | 2.5% | 0.4% |
| 16 | Pan-GATA6 | 1.7% | 2.5% | 24.6% | 9.6 % |
| | HA | 4.2% | 2.9% | 5.3% | 0.4% |
| 25 | Pan-GATA6 | 4.2% | 5.4% | 43.3% | 19.2 % |
| | HA | 7.1% | 5.0% | 22.5% | 2.9% |
